# An improved rhythmicity analysis method using Gaussian Processes detects cell-density dependent circadian oscillations in stem cells

**DOI:** 10.1101/2023.03.21.533651

**Authors:** Shabnam Sahay, Shishir Adhikari, Sahand Hormoz, Shaon Chakrabarti

## Abstract

Detecting oscillations in time series remains a challenging problem even after decades of research. In chronobiology, rhythms in time series (for instance gene expression, eclosion, egg-laying and feeding) datasets tend to be low amplitude, display large variations amongst replicates, and often exhibit varying peak-to-peak distances (non-stationarity). Most currently available rhythm detection methods are not specifically designed to handle such datasets. Here we introduce a new method, ODeGP (**O**scillation **De**tection using **G**aussian **P**rocesses), which combines Gaussian Process (GP) regression with Bayesian inference to provide a flexible approach to the problem. Besides naturally incorporating measurement errors and non-uniformly sampled data, ODeGP uses a recently developed kernel to improve detection of non-stationary waveforms. An additional advantage is that by using Bayes factors instead of p-values, ODeGP models both the null (non-rhythmic) and the alternative (rhythmic) hypotheses. Using a variety of synthetic datasets we first demonstrate that ODeGP almost always outperforms eight commonly used methods in detecting stationary as well as non-stationary oscillations. Next, on analyzing existing qPCR datasets that exhibit low amplitude and noisy oscillations, we demonstrate that our method is more sensitive compared to the existing methods at detecting weak oscillations. Finally, we generate new qPCR time-series datasets on pluripotent mouse embryonic stem cells, which are expected to exhibit no oscillations of the core circadian clock genes. Surprisingly, we discover using ODeGP that increasing cell density can result in the rapid generation of oscillations in the *Bmal1* gene, thus highlighting our method’s ability to discover unexpected patterns. In its current implementation, ODeGP (available as an R package) is meant only for analyzing single or a few time-trajectories, not genome-wide datasets.

## Introduction

From the rapid ultradian oscillations of p53, NF-*κ*B, Hes7, and the embryonic segmentation clock, to the slower seasonal flowering patterns in plants - oscillations in biological systems are ubiquitous across many length and time scales [1, 2]. These patterns, where repeatedly occurring peaks can be observed in the time series of interest, are often noisy, exhibit low amplitudes (peak- to-trough distance), and are non-stationary (peak-to-peak distance varies with time). Classic examples of such noisy oscillations can be observed in circadian clock gene expression [3, 4], feeding, eclosion and egg-laying rhythms [5, 6, 7]. This makes it hard to distinguish rhythmic patterns from noise, necessitating the development of quantitative approaches to accurately infer the existence of oscillations, and subsequently extract parameters such as time period and amplitude [8]. Developing principled approaches to detecting oscillations is also important in differential rhythmicity analyses, where genes can lose or gain rhythmicity after perturbations [9, 10, 11]. Finally, biological oscillators are often coupled, and investigating the nature and consequences of coupling often depends on the ability to carefully measure and detect the individual oscillations in the first place [12, 13, 14].

Over the last few decades numerous methods have been developed to address the oscillation detection problem arising from experiments which generate, for example, qPCR, RNA-seq, feeding, eclosion and egg-laying time series datasets. Existing non-parametric methods used for oscillation detection include JTK Cycle [15] and RAIN [16]. eJTK [17] is a more recently developed algorithm that improves on JTK Cycle by including non-sinusoidal reference waveforms. MetaCycle [18], an R package developed to identify oscillations, integrates the results of JTK Cycle, the Lomb Scargle Periodogram [19] (a parametric method) and ARSER [20] (a parametric method using autoregressive models, which cannot work on unevenly sampled data). Another existing R package for oscillation detection is DiscoRhythm [21], which builds on MetaCycle by additionally using the Cosinor [22] method (also parametric). Among the non-parametric methods, RAIN requires that the period to test for (or a range of periods) be specified beforehand when trying to identify oscillations. Similarly, JTK Cycle and MetaCycle also require information about the expected oscillation time period to be provided as inputs to the algorithm. More recently, neural network based approaches have been used for classifying oscillatory versus non-oscillatory datasets [23], but these do not allow learning of the full waveform, besides requiring a lot of training data to perform well. Finally, wavelet-based techniques [24, 25] can extract time-dependent phase and periods from temporal datasets, but are not designed to specifically detect oscillations or handle replicate data. More details and comparisons of a variety of oscillation detection algorithms can be found elsewhere - [8] provides a comprehensive review of earlier techniques while comparisons of more recent methods can be found in [26, 27, 28, 29, 30].

While these methods have improved over time and have become increasingly powerful, a number of challenges in oscillation detection are yet to be overcome [30] - (1) data is often available only at non-evenly spaced time points, which can make oscillation detection difficult, particularly with Fourier Transform based methods, (2) large error-bars at each time point from replicates are often not easy to incorporate into existing methods, (3) biological oscillations tend to be nonstationary (peak-to-peak distance varies over time), which parametric models cannot handle well due to difficulty in defining functional forms for such data, and finally (4) most current methods rely on calculating a p-value to classify a dataset as rhythmic versus non-rhythmic (for an exception, see [31]), but a major issue with p-value based approaches is that they model only the null, but not the alternative hypothesis [30, 32, 33, 34]. Existing methods often overcome one or few of these challenges, but no single method exists that addresses all these problems in a comprehensive manner.

To solve the above four challenges in a unified framework, here we develop ODeGP (**O**scillation **De**tection using **G**aussian **P**rocesses), a new approach to the oscillation detection problem combining Gaussian Process (GP) regression [35] with Bayesian model selection. Conceptually, the non-parametric nature of GPs allows ODeGP to flexibly model both stationary and nonstationary datasets, while the Bayesian model selection approach using Bayes factors allows us to model both the null as well as alternate hypotheses, unlike p-value based methods. In particular, we use a recently developed non-stationary kernel that allows us to model non-stationary datasets [36, 37], improving the accuracy of oscillation detection over many of the popular existing methods such as eJTK and MetaCycle. GPs also overcome issues related to unevenly spaced time-series data and can naturally incorporate error bars generated by replicates. Additionally, ODeGP provides a simple Bonferroni-type multiple hypothesis correction [38], though this approach currently limits its use to settings where only one or a few trajectories are to be analyzed, not genome-wide datasets. Finally, a major additional advantage of GPs is that they provide the ability to quantify uncertainty predictions at any time point, including test points where no experimental data has been collected [35].

We extensively compare the performance of ODeGP with eight other existing methods on both simulated and experimental datasets and demonstrate that it is consistently better and more sensitive at identifying oscillations. We also find that the Bayes factor usually has a large separation between oscillatory and non-oscillatory experimental datasets, suggesting that it is a good metric for the classification problem. Finally, to test the usefulness of ODeGP in learning patterns in new datasets, we generate time-series circadian clock gene expression profiles using mouse embryonic stem cells (mESCs). Intriguingly, we find that oscillations of *Bmal1* can be induced within a few days in mESCs by increasing cell density and that these oscillations get suppressed with the addition of MEK/ERK and GSK3b inhibitors. This interesting result adds to previous observations that while pluripotent mESCs exhibit no clock gene oscillations, retinoic acid mediated differentiation can induce oscillations after about two weeks [39, 40]. Our results indicate that increasing cell density might mimic the effects of directed differentiation, but with faster kinetics of emergence of circadian gene oscillations.

## Methods

### ODeGP - Gaussian Process regression for oscillation detection

A brief mathematical introduction to the theory of Gaussian Processes (GPs) is provided in SI Section 3. Here we provide the basic outline of our oscillation detection algorithm using GPs. The time points at which gene expression data is collected using qPCR are specified as a list *X*. If the data for *d* distinct replicates is available for the times given in *X*, these *d* lists *Y*_1_, *Y*_2_, …, *Y*_*d*_ are taken as input. Otherwise, if the averaged qPCR data *Y* is available along with the corresponding error bar values *S* for each time point, *Y* and *S* are taken as input. In either case, the qPCR data may be collected at irregular intervals or have missing points. In the former case, *Y* and *S* are calculated from *Y*_1_, *Y*_2_, …, *Y*_*d*_ before proceeding. *Y* is then detrended via linear regression to remove long-term trends, and zero-centred to simplify the computations involved in performing Gaussian process regression.

The entries of the covariance matrix Σ_*XX*_ are determined by a positive semi-definite kernel function *K*. ODeGP uses two different kernel functions, *K*_*D*_ and *K*_NS_. The diagonal kernel *K*_*D*_(*x, x*^′^) = *ϵ*^2^ *· δ*_*xx*′_ is used to represent non-oscillatory functions, where *δ* is the Kronecker delta. This kernel encodes the prior belief that there is no correlation between the values of the data at different time points, thus all the non-diagonal entries of the covariance matrix are 0. To represent oscillatory functions, the non-stationary kernel *K*_NS_(*x, x*^′^) = *w*(*x*)*w*(*x*^′^)*k*_gibbs_(*x, x*^′^) cos(2*π*(*xμ*(*x*) *− x*^′^*μ*(*x*^′^)) + *ϵ*^2^ *· δ*_*xx*′_ is used, where *k*_gibbs_ is the Gibbs kernel. The hyperparameters *w* and *μ*, along with *l* (which is part of the expression of *k*_gibbs_), are functions of *x*. Thus the covariance matrix will depend on *x* and is not solely a function of |*x − x*^′^|.

The error bar values *S* are incorporated into the covariance matrices generated by these kernels as follows: for 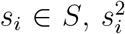 is added to the *i*th diagonal entry of the matrix. These terms model local noise, which arise from technical variations amongst the replicates. Global noise, which includes other sources of variations beyond technical noise, is modeled by a hyperparameter *ϵ* such that *ϵ*^2^ is added to all diagonal entries of the covariance matrix. This *ϵ*^2^ term is included in the expressions of *K*_*D*_ and *K*_*NS*_, allowing us to learn the best value of *ϵ* along with the remaining hyperparameters.

Optimal hyperparameters for the non-stationary and diagonal kernels are obtained through maximization of the marginal log-likelihoods (MLL) (see Equation 17 in the SI), to obtain the optimal 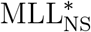 and 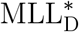 respectively. The kernel that represents a better prior belief for the given dataset is identified through Bayesian model selection. The Bayes factor is defined as the ratio of the marginal likelihood of two competing hypotheses. Here, it is calculated as 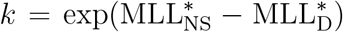 and compared to a decided threshold *T*. If *k ≥ T*, the dataset is declared to be oscillatory, otherwise not. A discussion on the choice of *T* is provided in the Discussion section.

Finally, while ODeGP has been designed primarily to detect rhythms in single time-series trajectories, it also provides a Bonferroni-type multiple-hypothesis correction term if multiple trajectories from the same dataset are intended to be analyzed simultaneously. A single multiplicative correction term to the Bayes Factors (which acts as the prior odds) is calculated based on the number of trajectories to be analyzed: 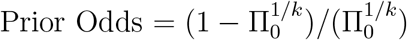, where *k* is the number of trajectories and Π_0_ is by default taken to be 0.5. The prior odds, multiplied to each of the Bayes Factors of the different trajectories analyzed, provides the posterior odds that can be used to judge rhythmicity in the multiple hypothesis setting. More details are provided in Section 4 of the SI. The use of this Bonferroni-type correction factor currently limits ODeGP to the analysis of one or few trajectories only, as opposed to genome-wide datasets, since the Bonferroni correction is very stringent and greatly increases false negatives when many comparisons are performed.

### Cell lines and culture conditions

A passage 12 (p12) mESC line (E14TG2a.4) was expanded under conditions expected to maintain pluripotency, and all cells used for the experimental data reported here were within 10 additional passages. The p12 cells were thawed and expanded on 0.1% gelatin-coated, cell-culture treated 10 cm plastic dishes. GTES ES cell media (GMEM, 15% FBS, Glutamax, Sodium Pyruvate, Non-Essential Amino Acids, BME, and LIF) was used to propagate the cells in pluripotent conditions. Cells were passaged at approximately 70% confluency to avoid crowding-induced differentiation. For all experiments that required thawing out new vials, the cells were always initially maintained in cell-culture treated plastic dishes coated with 0.1% gelatin, and then transferred to various other conditions, such as glass dishes with fibronectin or laminin coating.

### qPCR experiments - data collection protocol and error analysis

For the qPCR experiments, cells were collected in Trizol, total RNA was extracted and converted to cDNA, and finally, qPCR was performed using the SYBR Green dye. Each time point (beginning from time 0, corresponding to immediately after Dex synchronization) corresponds to cells obtained from an independent well of a 24-well plate. To avoid any artifacts in cell synchronization due to differences in cell density at different times of cell collection, we devised a protocol to ensure an approximately equal number of cells collected for every time point: instead of synchronizing all the samples at one time point and collecting cells at different time points, we synchronized cells at different times and Trizol collected the cells at a single time. Details of the protocol, seeding densities, and various substrate conditions for the samples are provided in SI Section 1. The error analysis is explained in SI Section 2.

## Results

### ODeGP - an oscillation detection algorithm based on Gaussian Processes

We developed a new method for detecting oscillations, ODeGP (**O**scillation **De**tection using **G**aussian **P**rocesses), combining Gaussian Process (GP) regression and Bayesian inference. A brief intuition of how GP regression works is provided in Figure 1A and an outline of the ODeGP pipeline is displayed in Figure 1B. In brief, GP regression is a non-parametric approach to learning non-linear trends in data, where instead of specifying a function and learning its optimal parameters (parametric regression), the functional form itself is learnt by specifying a prior over functions [35]. Placing a prior over functions is achieved by the use of a multivariate Gaussian (Figure 1A, left), whose covariance matrix is determined by a kernel that prioritizes certain classes of functions based on prior expectations of smoothness and characteristic length-scales associated with the problem of interest. After data is obtained, the instantiations of the prior that best describe the data are obtained via the posterior (Figure 1A, right), which can be described via the posterior mean and variance. More details can be found in Methods and in the SI Sections 3 and 4.

**Figure 1.**
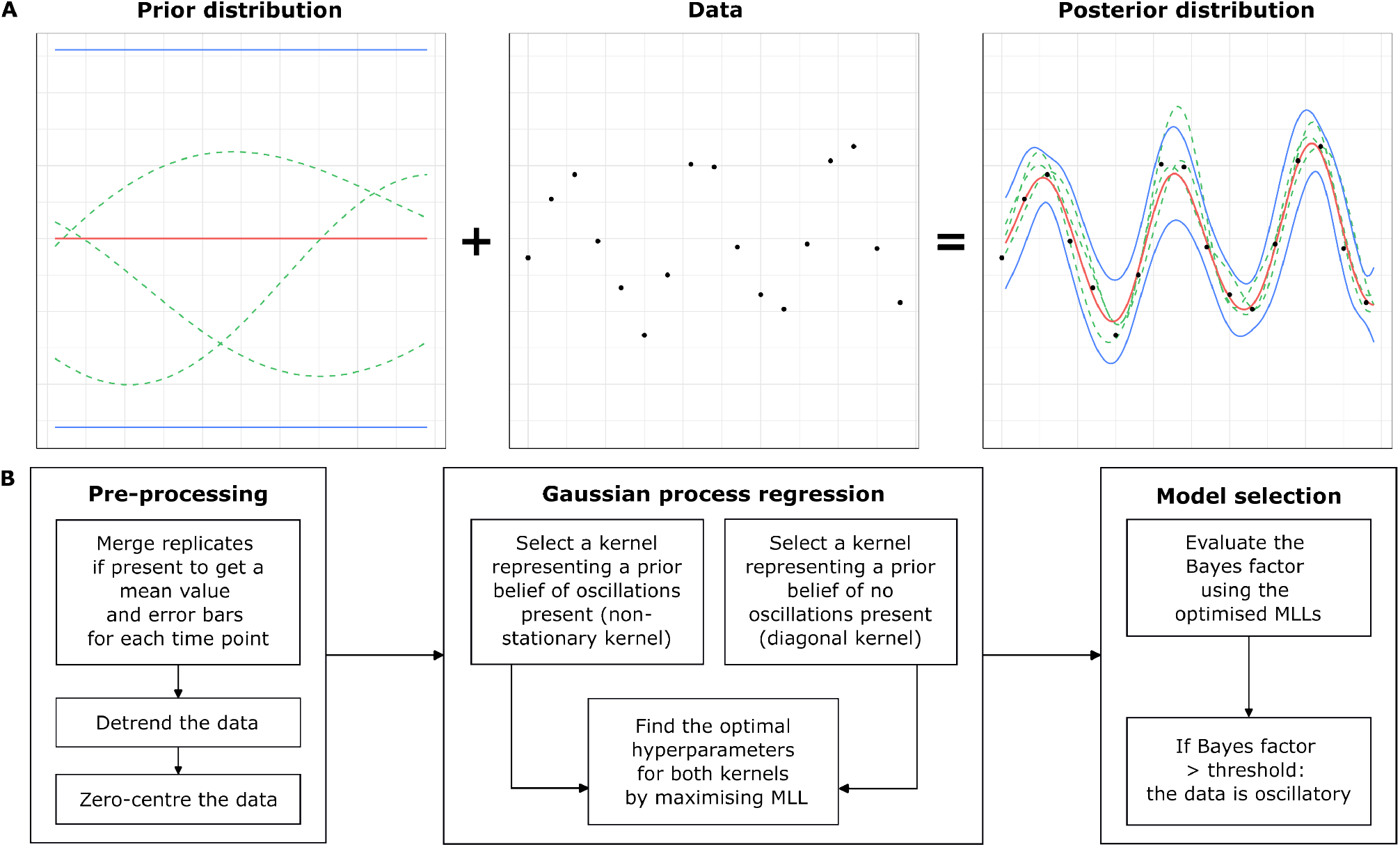
An overview of the workflow of ODeGP. (A) An intuitive schematic of Gaussian Process (GP) regression, where a GP is an indexed collection of random variables with a multivariate normal joint probability density. Left: A GP prior defining a distribution over functions (instantiations of the functions are shown in green lines; mean and confidence intervals are in red and blue respectively). Middle: Observed data. Combining the observed data and the GP prior allows the construction of the data likelihood. Right: Posterior distribution generated after Bayesian inference, which models the given data closely. Green lines are instantiations of the posterior, red and blue lines are the posterior mean and standard deviation respectively. (B) The workflow of ODeGP. The pre-processing stage involves formatting and normalizing the data. Next, two separate regressions are performed with the diagonal and non-stationary kernels respectively. The optimized marginal log likelihood (MLL) of the data is found for each case. These MLL values are then used to compute the Bayes factor for model selection. This Bayes factor is the final output metric used to determine whether the data is oscillatory or not.

ODeGP detects oscillations in time-series data by initializing two GP models or kernels (one encoding a belief of the data containing oscillations, and the other encoding a belief of oscillations being absent), performing GP regression on the data with each kernel separately by optimizing their respective marginal log-likelihoods (MLLs), and finally comparing these likelihoods to determine the model that better describes the data. Optimization of the MLL automatically incorporates the trade-off between maximizing the fitting of the data while minimizing model complexity. This complexity is represented by the determinant of the covariance matrix in the expression for the MLL (Equation 17 in the SI). This term can penalize an increase in the number of kernel hyperparameters used to compute the covariance matrix, and thus prevent overfitting of the data. A more detailed analysis of this is presented in SI Section 6, Figure S1 and Table S7. Finally, the possibility of performing corrections for multiple hypothesis testing is also provided - for details, see Methods and SI Section 4.

We generated a wide variety of simulated datasets where the ground truth (oscillatory or non-oscillatory) is known. These datasets are summarized in Table 1, and comparisons of various methods on these datasets are discussed in the next sections (see also SI Section 5). To mimic experimental qPCR data, all simulated waves were generated as sets of 3 replicates, with a total duration of 48 hours and an interval of 3 hours between consecutive observations. Two relative levels of noise (low and high), along with two different fractions of missing data (none and half), were used to create further variation among the datasets. Finally, three different ways of generating non-stationary data were tested (details in Section 5 of the SI).

**Table 1:** Categories of simulated data used for comparing the performance of existing oscillation-detection methods with ODeGP.

| All datasets: 3 replicates, 48 hr duration, 3 hr spacing,<br>Noise level: low/high, Fraction of missing data: 0/0.5 |  |  |
| --- | --- | --- |
| Simulated non-oscillatory datasets | Gaussian noise (Random sample from $\mathcal{N}(0, \sigma)$ at each timepoint) | |
| Simulated oscillatory datasets | Stationary symmetric data (combinations of sine waves) | Stationary asymmetric data (sawtooth waves) |
|  | Non-stationary symmetric data (continuously decreasing or randomly varying time period) | Non-stationary asymmetric data (randomly varying time period) |

### Detecting oscillations in simulated stationary datasets

We first tested ODeGP on simulated stationary data. Datasets consisting of both non-oscillatory and stationary oscillatory waves were generated, and the ability of each method to distinguish the two was evaluated through the construction of receiver operating characteristic (ROC) curves.

Non-oscillatory waves were simulated by randomly sampling from a standard normal distribution with standard deviation *Σ* at each timepoint in consideration: *f* (*t*) = *𝒩* (0, *Σ*), as shown in Figure 2A. The three replicates are shown with red, blue and green lines. Symmetric stationary waves, as in Figure 2B, were generated by the addition of two sine waves: *f* (*t*) = *A*_1_sin(2*πt/τ*_1_) + *A*_2_sin(2*πt/τ*_2_) + *𝒩* (0, *Σ*).

**Figure 2.**
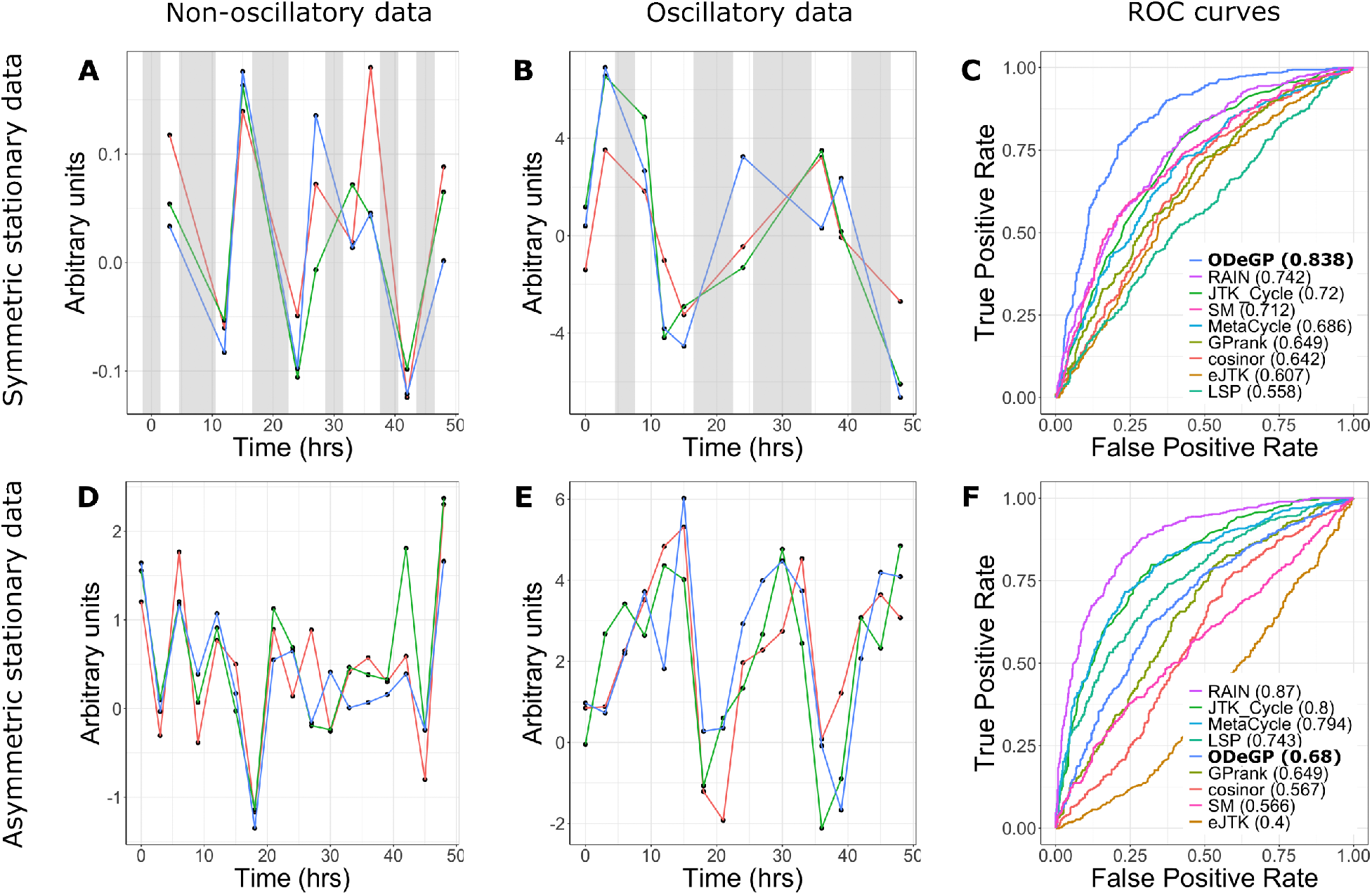
Detecting oscillations in simulated stationary datasets. All waves shown are sets of three replicates. (A) Simulated non-oscillatory data with *Σ* = 0.1 and half the points missing. (B) Simulated symmetric stationary data with *A*_1_ = *A*_2_ = 3, *τ*_1_ = 18, *τ*_2_ = 26, *Σ* = 0.1 and half the points missing. (C) ROC curves for all methods considered on the dataset consisting of waves generated like (A) and (B). Numbers in brackets denote AUC values. (D) Simulated non-oscillatory data with *Σ* = 1 and no points missing. (E) Simulated asymmetric stationary data with *A* = 5, *τ* = 18 and *Σ* = 1. (F) ROC curves for all methods considered on the dataset consisting of waves generated like (D) and (E). Grey shaded areas represent regions with missing data. LSP - Lomb Scargle Periodogram; SM - spectral mixture kernel.

The performance of various methods in terms of correctly identifying the presence of oscillations in symmetric stationary waves was evaluated by generating ROC curves on a collection of 500 oscillatory and 500 non-oscillatory waves. These ROC curves for data corresponding to the waves in Figure 2A and 2B are shown in Figure 2C. ODeGP performs the best on this set of 1000 waves, with an AUC value of 0.838. AUC values for all other datasets are reported in SI Section 5.

Similarly, performance on asymmetric stationary oscillatory waves was evaluated by generating ROC curves on a collection of 500 non-oscillatory waves (as shown in figure 2D) and 500 asymmetric stationary oscillatory waves. The latter were generated using the functional form of sawtooth waves: 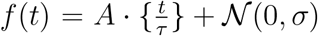, where *{}* represents the fractional part function.

Figure 2E shows a wave generated as such. RAIN performs the best on this set of 1000 waves as seen in Figure 2F with an AUC value of 0.87, while the non-stationary kernel has a relatively poorer AUC value of 0.68.

The relative performance of the methods tested on all stationary datasets generated is summarised in Table 2 (all AUC values are provided in SI Section 5). ODeGP is consistently among the best 3 performing methods in all symmetric stationary datasets considered. Cosinor and eJTK, which are among the best 3 methods the second-most times, also fall among the worst 3 methods a significant number of times. In asymmetric stationary datasets, RAIN is among the best 3 methods the most consistently, whereas ODeGP here has a relatively average performance (neither being many times among the best 3 nor many among the worst 3). RAIN’s better performance compared with our method on these datasets is expected because it separately groups the rising and falling parts of the waves for comparison, boosting its ability to identify asymmetric waveforms [16].

**Table 2:** Comparison of the number of times each method tested appears among the best and worst performing 3 methods, in all stationary datasets considered. LSP - Lomb Scargle Periodogram; SM - spectral mixture.

| Stationary Datasets |  |  |  |  |
| --- | --- | --- | --- | --- |
| Method | Symmetric Datasets (Total: 9) |  | Asymmetric Datasets (Total: 8) |  |
|  | Times among<br>best 3 methods | Times among<br>worst 3 methods | Times among<br>best 3 methods | Times among<br>worst 3 methods |
| Cosinor | 5 | 3 | 3 | 4 |
| eJTK | 5 | 4 | 4 | 4 |
| GPrank | 0 | 6 | 0 | 5 |
| JTK_Cycle | 1 | 0 | 3 | 0 |
| LSP | 0 | 6 | 1 | 4 |
| MetaCycle | 1 | 0 | 4 | 0 |
| ODeGP | 9 | 0 | 2 | 1 |
| SM Kernel | 2 | 5 | 0 | 6 |
| RAIN | 4 | 3 | 7 | 0 |

### Detecting oscillations in simulated non-stationary datasets

Since experimental time-series qPCR data tends to be non-stationary (peak-to-peak distance varies with time), we next tested the ability of each method to distinguish non-stationary oscillatory waves from non-oscillatory waves.

Symmetric non-stationary oscillatory waves with a monotonically decreasing time period, as shown in Figure 3B, were generated using the functional form 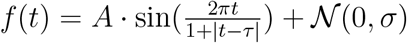, where *τ ≥* 48. The ROC curves in Figure 3C indicate that ODeGP performs the best on a dataset consisting of 500 non-oscillatory waves generated like in Figure 3A and 500 symmetric non-stationary oscillatory waves generated like in Figure 3B, with an AUC value of 0.814.

**Figure 3.**
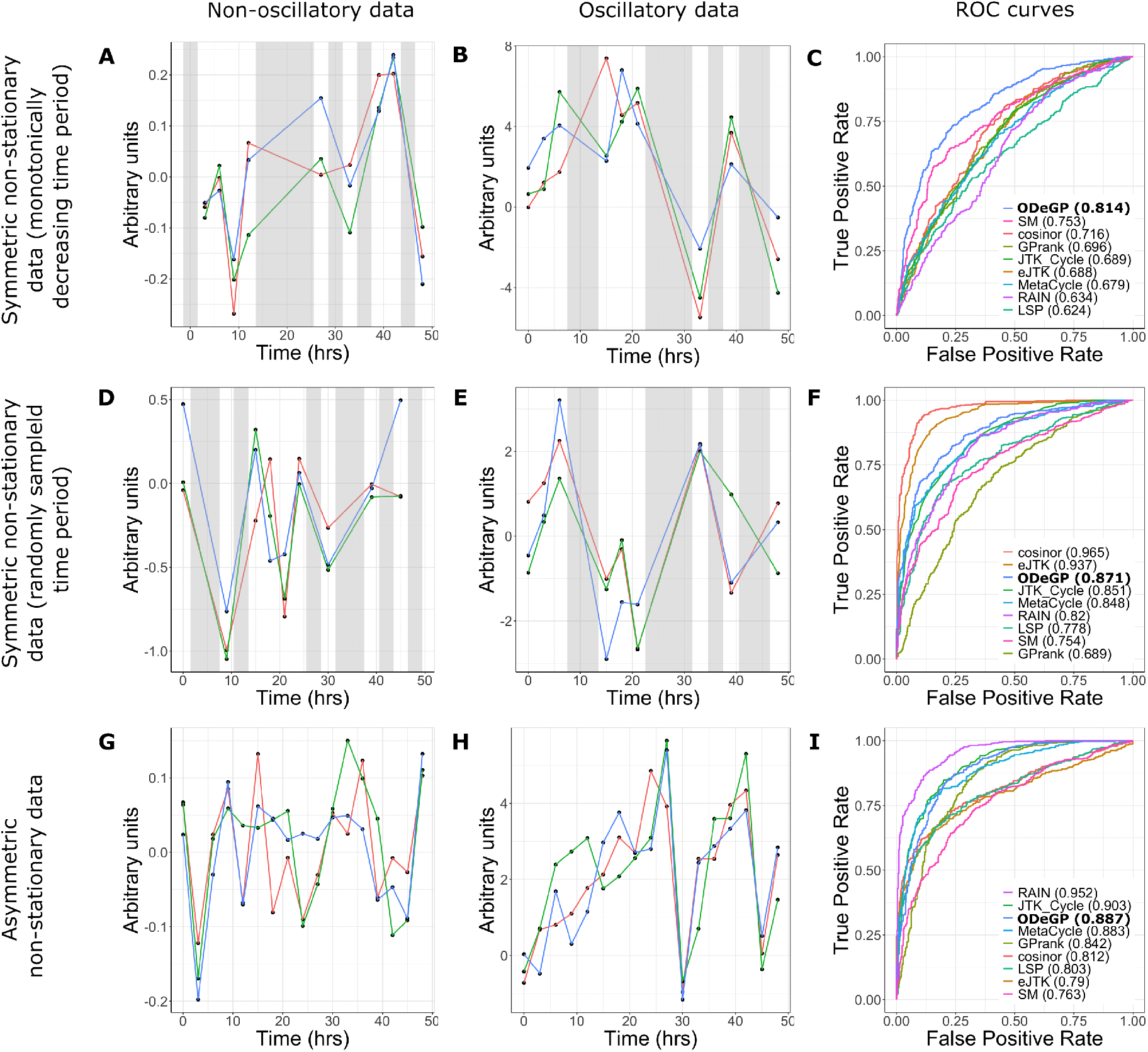
Detecting oscillations in simulated non-stationary datasets. All waves shown are sets of three replicates. (A) Simulated non-oscillatory data with *Σ* = 0.1 and half the points missing. (B) Simulated symmetric non-stationary data with a monotonically decreasing time period, with *A* = 5, *τ* = 72, *Σ* = 0.1 and half the points missing. (C) ROC curves for all methods tested on the dataset consisting of waves generated like (A) and (B). (D) Simulated non-oscillatory data with *Σ* = 0.5 and half the points missing. (E) Simulated symmetric non-stationary data with a randomly varying time period, with *A* = 1.5, *μ* = 24, *Σ* = 1.33, *Σ* = 0.5, and half the points missing. (F) ROC curves for all methods tested on the dataset consisting of waves generated like (D) and (E). (G) Simulated non-oscillatory data with *Σ* = 0.1 and no points missing. (H) Simulated asymmetric non-stationary data with *A* = 5, *τ*_1_ = 12, *τ*_2_ = 30, and *Σ* = 0.1. (I) ROC curves for all methods tested on the dataset consisting of waves generated like (G) and (H). Grey shaded areas represent regions with missing data. Numbers in brackets in panels (C), (F) and (I) correspond to AUC values. LSP - Lomb Scargle Periodogram; SM - spectral mixture kernel.

Since there can be many ways of generating non-stationary data, an additional approach to generating time-varying periodicities in a single wave was explored. Symmetric oscillatory waves were generated with a randomly varying time period instead of a monotonically decreasing one, as follows: at the start of each new oscillation, a time period *τ* is sampled from *N* (*μ, Σ*). If the total time elapsed up to the start of this new oscillation is *ϕ*, then 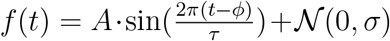 for *ϕ ≤ t < ϕ* + *τ*. Figure 3E shows a wave generated in this way. Figure 3F demonstrates that cosinor is the best performing method on the dataset consisting of 500 non-oscillatory waves generated like in Figure 3D and 500 symmetric non-stationary oscillatory waves generated like in Figure 3E. While ODeGP has a poorer AUC value of 0.871 compared to 0.965 for Cosinor and 0.937 for eJTK, it still remains within the top three performing methods.

Asymmetric non-stationary oscillatory waves (like shown in Figure 3H) were generated using a similar principle, but with a sawtooth waveform instead of a sine waveform. At the start of each new oscillation, a time period *τ* is sampled from *U* (*τ*_1_, *τ*_2_). If the total time elapsed up to the start of this new oscillation is *ϕ*, then 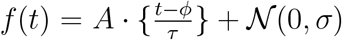 for *ϕ ≤ t < ϕ* + *τ*, where *{}* represents the fractional part function. RAIN can be seen as the best performing method in Figure 3I for the dataset consisting of 500 non-oscillatory waves generated like in Figure 3G and 500 asymmetric non-stationary oscillatory waves generated like in Figure 3H. ODeGP however remains among the top three methods here as well.

A comparison of the performance of the methods tested across all simulated non-stationary datasets is shown in Table 3 (AUC values from all datasets are provided in SI Section 5). ODeGP is the best performer for symmetric non-stationary datasets, being among the top 3 methods (in terms of AUC values) for all 15 datasets tested. In the non-stationary case as well, RAIN again emerges as the best method for asymmetric datasets. ODeGP shows a slight improvement in its relative performance on non-stationary datasets compared with stationary asymmetric datasets, being among the best 3 methods for a larger fraction of the datasets, and also never falling among the worst 3 methods.

**Table 3:** Comparison of the number of times each method tested appears among the best and worst performing 3 methods, in all non-stationary datasets considered. LSP - Lomb Scargle Periodogram; SM - spectral mixture

| Non-stationary Datasets |  |  |  |  |
| --- | --- | --- | --- | --- |
| Method | Symmetric Datasets (Total: 15) |  | Asymmetric Datasets (Total: 12) |  |
|  | Times among best 3 methods | Times among worst 3 methods | Times among best 3 methods | Times among worst 3 methods |
| Cosinor | 9 | 2 | 4 | 5 |
| eJTK | 7 | 4 | 4 | 6 |
| GPrank | 2 | 4 | 0 | 6 |
| JTK_Cycle | 4 | 2 | 8 | 0 |
| LSP | 0 | 14 | 0 | 8 |
| MetaCycle | 2 | 6 | 5 | 1 |
| ODeGP | 15 | 0 | 5 | 0 |
| SM Kernel | 4 | 7 | 0 | 10 |
| RAIN | 2 | 6 | 10 | 0 |

### ODeGP provides increased sensitivity at distinguishing oscillatory vs non-oscillatory patterns in noisy qPCR datasets

After analyzing simulated datasets, we next evaluated ODeGP’s ability to distinguish oscillatory and non-oscillatory patterns in experimental datasets and benchmarked its performance against existing methods. We started with a published dataset on primary mouse marrow stromal cells, where the expression levels of a number of circadian clock genes were measured over 48 hours using qPCR [4]. The cells were either treated with Dexamethasone (Dex) which is expected to synchronize or stimulate clock gene expression oscillations, or with vehicle (DMSO) where oscillations are not expected. Three independent experiments were done at each time point, thereby providing error bars for the expression levels as well. This dataset, therefore, represented a good test case for applying our rhythm detection method, since the ground truth is known.

The results from our analysis of the *Rev-ERBβ* and *Per1* genes are shown in Figure 4 (other genes are shown in Figure S2, SI Section 6). Raw data for *Rev-ERBβ*, which exhibited large amplitude oscillations, are shown in Figure 4A and 4B along with the GP posteriors (mean and standard deviation) generated using the non-stationary kernel. Besides an AUC of its ROC curve being close to one, an additional characteristic of a good binary classifier is its ability to produce an output metric that is well-separated for the two classes in consideration - in our case, non-oscillatory and oscillatory. Applying ODeGP on the vehicle treated data (Figure 4A) produces a Bayes factor of 16.50, whereas it produces a Bayes factor of 50133.83 on the synchronized data in Figure 4B, a separation of more than three orders of magnitude. The significant separation between these two values shows that ODeGP is able to make a clear distinction between the non-oscillatory (or weakly oscillatory; see more in the Discussion section) and oscillatory qPCR data. The same trend is observed in the other genes we analyzed (Figure S2), where there is at least an order of magnitude increase in the Bayes Factor, usually even more, when the data is oscillatory.

**Figure 4.**
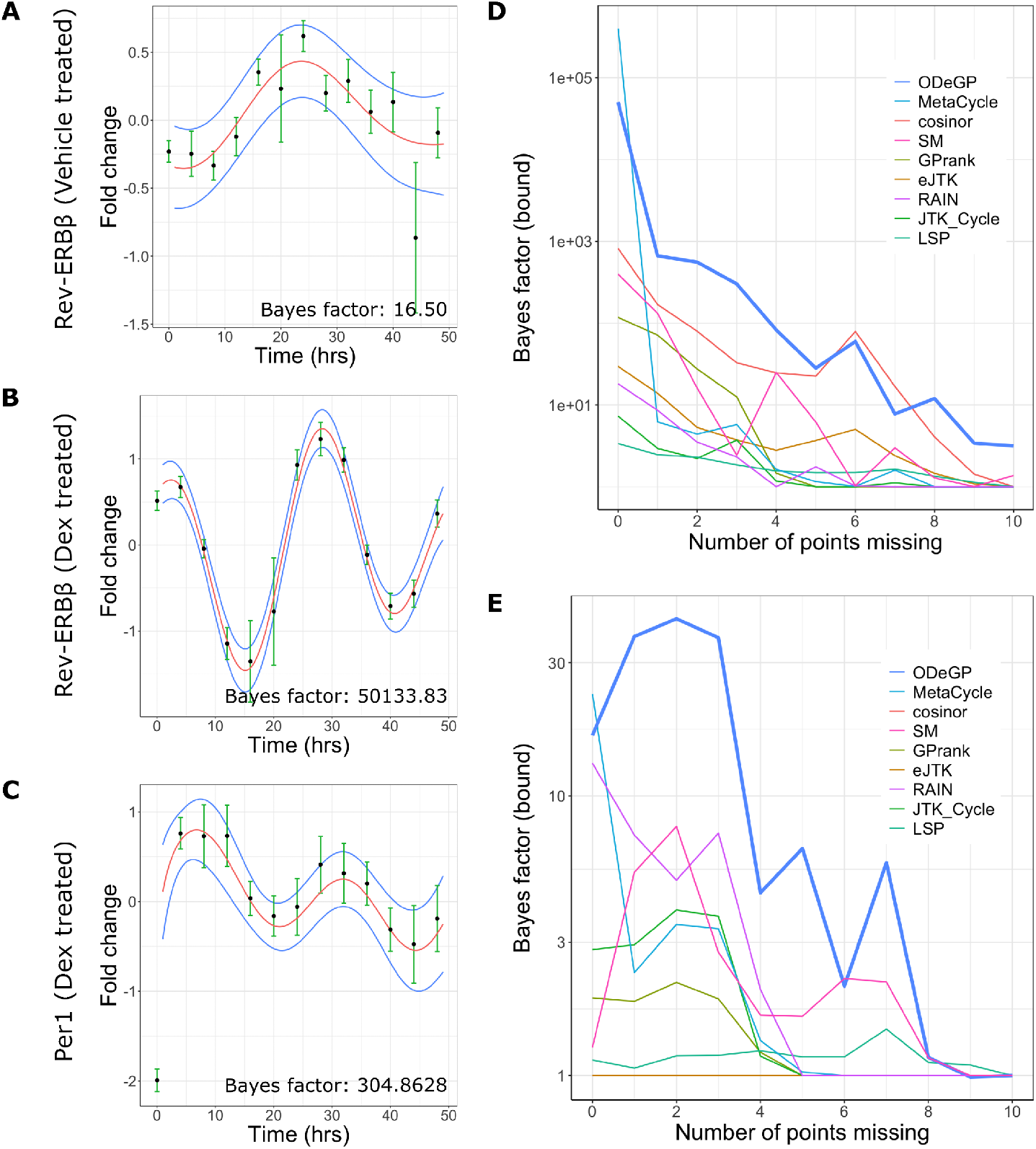
Application of ODeGP on *Rev-ERBβ* and *Per1* expression data [4] demonstrates the sensitivity of ODeGP at distinguishing oscillatory from non-oscillatory data. (A) The relative gene expression of *Rev-ERBβ* when treated with DMSO, measured at 4-hour intervals over a 48-hour duration, is shown by the black points with green error bars indicating an average taken over 3 biological replicates. Defining a GP prior using the non-stationary kernel and performing GP regression on this data produces the posterior distribution shown (mean in red, standard deviation in blue). (B) Relative gene expression of *Rev-ERBβ* when treated with Dex, representing an oscillatory dataset, is shown by the black points with green error bars. The GP posterior of the non-stationary kernel applied to this data is shown with the posterior mean and standard deviation in red, and blue respectively. (C) Relative gene expression of *Per1*, when treated with Dex, is shown by the black points with green error bars. This dataset represents ground-truth oscillatory data, but with smaller amplitude oscillations as compared to (B). The GP posterior of the non-stationary kernel applied to this data is shown with the posterior mean and standard deviation in red, and blue respectively. ODeGP classifies this as oscillatory with much more confidence than other existing methods (see main text and Table 4). (D) Points from the raw data in (B) were removed one by one in a random order, and all methods were applied to the resulting downsampled data at each step. The variation of the resulting Bayes factor (bounds) with increasing number of missing points is shown for each method. The thick blue line represents the ODeGP Bayes factor. (E) Analysis similar to panel (D) but for the raw data in panel (A).

The appropriateness of the non-stationary kernel for the classification problem is also highlighted by the narrower confidence intervals of the GP posterior in Figure 4B compared to those in Figure 4A, which indicates that the non-stationary model is a less complex model for the oscillatory data than for the non-oscillatory data. Similar separations of Bayes factors and narrower confidence intervals were observed when applying ODeGP on the corresponding data for the *Per2* and *Npas2* genes as well (Fig. S2).

**Table 4:** Comparison of output metrics of all methods tested on the low-amplitude *Per1* oscillation data shown in Figure 4C. p-values/Q-values were converted to corresponding Bayes factor bound [33] values where applicable to allow comparison with the Bayes factor returned by ODeGP. While eJTK, MetaCycle and RAIN are the only other methods that correctly classify the dataset as oscillatory (based on a p-value cutoff of 0.05), the large difference between ODeGP’s Bayes factor and the upper bounds generated by these methods suggest that ODeGP is more sensitive at detecting oscillations.

| Method | Metric type | Metric value | Bayes factor (bound) |
| --- | --- | --- | --- |
| Cosinor | Q-value | 0.08537 | 1.7512 |
| eJTK | p-value | 0.03577 | 3.0879 |
| GPrank | log of Bayes factor | -6.7185e-05 | 0.9999 |
| JTK_Cycle | p-value | 0.12445 | 1.4185 |
| LSP | p-value | 0.43067 | 1.0140 |
| MetaCycle | p-value | 0.04496 | 2.6376 |
| ODeGP | Bayes factor | 304.8628 | 304.8628 |
| SM Kernel | Bayes factor | 1.2632 | 1.2632 |
| RAIN | p-value | 0.00617 | 11.7191 |

We next asked if ODeGP is more sensitive at detecting oscillations compared to other existing methods. To evaluate this, we analyzed *Per1* data from the same paper [4], which visibly exhibited lower amplitude oscillations with larger error bars (Figure 4C). To compare the p (or Q) values generated by other methods against the Bayes factor produced by ODeGP, we converted the p-values to Bayes factor bounds [33] (Table 4). The Bayes factor bound represents an upper bound on Bayes factors corresponding to a given p-value, under very general assumptions on the alternative hypothesis [33]. As is clear from Table 4, the only methods besides ODeGP that correctly classified the data as rhythmic, were eJTK, MetaCycle and RAIN (p-values less than 0.05). However, the Bayes factor generated by ODeGP was much larger compared to the upper bound values of eJTK, Metacycle or RAIN, demonstrating that ODeGP correctly classified the oscillations with much more confidence. Indeed, if recent guidelines for rejecting the null based on p-values less than 0.005 ([33]; see Discussion section) were to be used, ODeGP would be the only method to correctly classify this dataset as oscillatory. The data’s oscillatory trend was also captured well in ODeGP’s non-stationary kernel posterior shown in Figure 4C.

To more systematically test ODeGP’s sensitivity of detecting oscillations against existing methods, we compared the Bayes factor generated by ODeGP with the Bayes factor bound values produced by the other methods across an increasingly down-sampled dataset. The raw data for Rev-ERB*β* in Dex-treated cells (Figure 4B), which showed large amplitude oscillations, was considered as a starting point. Points were then removed from this dataset one by one in a random order, and at each step, all methods were applied to the sub-sampled data. Figure 4D demonstrates the rapid decrease in the Bayes factor and Bayes factor bounds with increasing missing points. At zero points missing all methods perform well (i.e. produce large Bayes factor bound values), though ODeGP and MetaCycle are distinctly better. As the number of missing points increases, the ODeGP Bayes factor (blue thick line in Figure 4D) most consistently maintains a larger value compared to the Bayes factor bounds obtained from other methods. ODeGP thus performs better at identifying the data as oscillatory at an extent of missing points that causes other methods to fail. We also performed a similar downsampling analysis for the weakly oscillatory dataset in Figure 4A, the results of which are shown in Figure 4E. On comparing Figures 4D and E, it is evident that for most points on the x-axis, the strong versus weak oscillation Bayes Factors are best separated for ODeGP.

### Cell-density dependent rapid emergence of oscillations in mouse embryonic stem cells

Finally, we tested ODeGP’s ability to quantify oscillatory behaviour in new qPCR datasets, that could enable novel biological discoveries. For this purpose, we used early passage pluripotent mouse embryonic stem cells (mESCs). Previous work has demonstrated that pluripotent stem cells do not exhibit oscillations of the core circadian clock genes, even though the genes are expressed in these cells [39, 40]. We first confirmed these well-established results by culturing mESCs on a variety of substrates in the presence of LIF where pluripotency is expected to be maintained (gelatin on plastic, fibronectin on glass and laminin on glass; details in the SI Section 1). We synchronized cells with Dex and collected the cells over a period of 24 hours at intervals of 3 hours. Consistent with the previous literature [39, 40], application of our algorithm confirmed that there were no oscillations in any of the tested conditions, since the Bayes factors were in the range 2-8 (Figure 5A). As we had observed in the last section, in known cases where there are no oscillations, the Bayes factor tends to be of the order of 10 while the presence of real oscillations pushes up the Bayes factor by one or two orders of magnitude.

**Figure 5.**
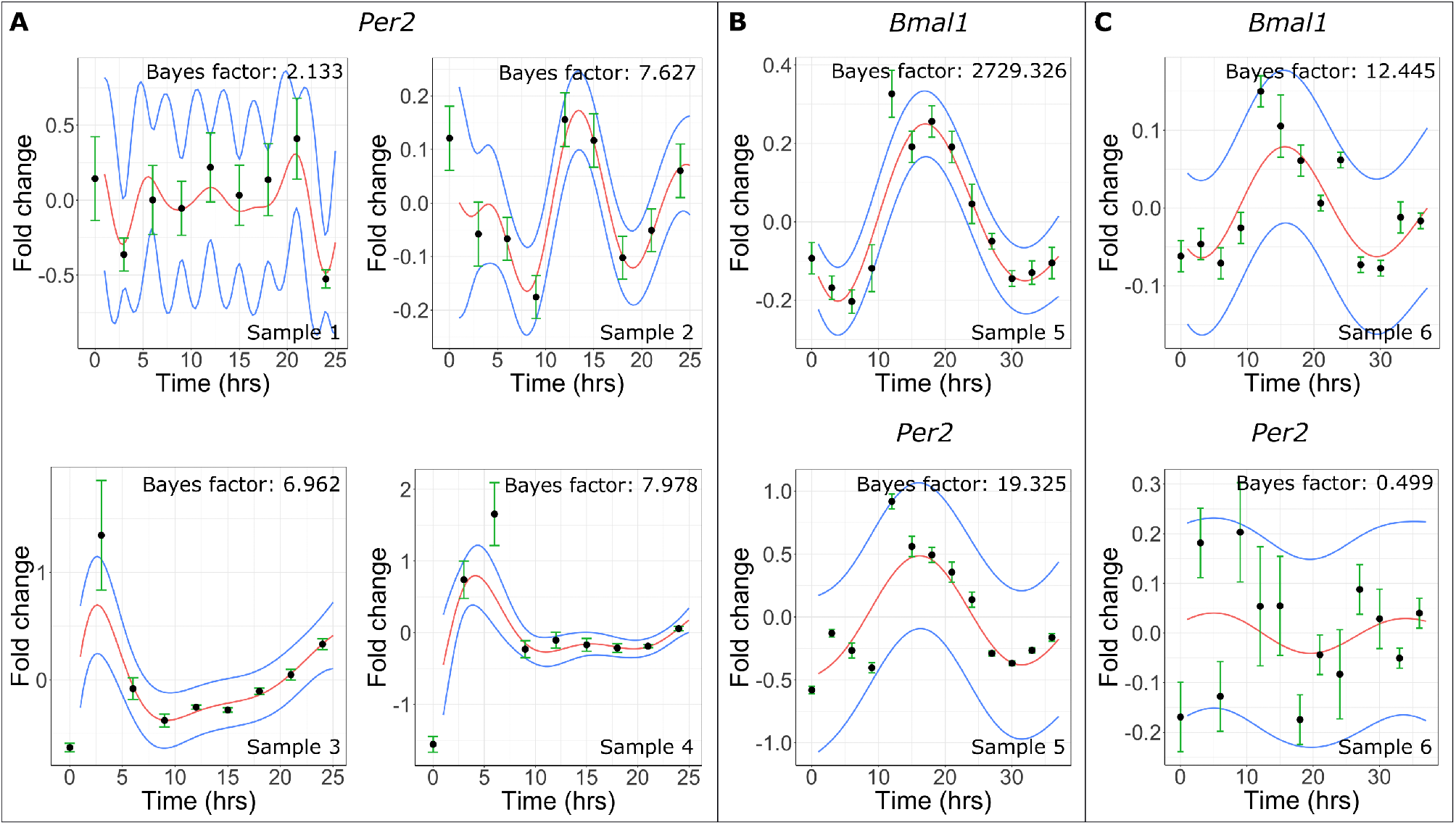
Cell-density dependent rapid emergence of oscillations in mESCs. GP posteriors of the non-stationary kernel (mean in red, standard deviation in blue) for (A) *Per2* gene expression from low-density samples 1,2,3 and 4 (see detailed descriptions in the SI), (B) *Bmal1* and *Per2* expression for high-density sample 5, and (C) *Bmal1* and *Per2* expression for high-density (but 2i treated) sample 6. In all plots, black circles represent the mean qPCR measurement from three technical replicates.

We next asked if increasing the cell density could lead to the generation of oscillations of circadian clock gene expression. Previous work has demonstrated that about two weeks of Retinoic Acid (RA) induced differentiation can induce oscillations in mESCs [39, 40], but to the best of our knowledge, the kinetics of oscillation development upon cell density increase has not been explored in this cell type. Since higher cell density can potentially cause some amount of differentiation in stem cells [41], we explored the consequences of doubling the number of cells in our culture dish before Trizol extraction and gene expression quantification. We also extended the time over which cells were collected from 24 to 36 hours, to allow for better detection of potential oscillations. Interestingly, though the higher cell density was maintained for only about 3-4 days (see SI Section 1 for details), we saw clear signs of oscillation in *Bmal1* (Bayes factor of 2729) and weak oscillations in *Per2* (Bayes factor of 19), as can be seen in Figure 5B. This was in contrast to the earlier studies using RA, where oscillations emerged only after two weeks [39]. To check whether cell differentiation could potentially have played a role in generation of these oscillations, we added the dual inhibitors of MEK/ERK and GSK3b (commonly called 2i) to an otherwise identical experimental set up with the higher cell density. 2i has been shown to differentially kill cells with low *Nanog* expression levels, which are prone to undergoing differentiation [42]. This time we found much lower Bayes factors for both *Bmal1* as well as *Per2* - 12.4 and 0.5 respectively (Figure 5C), clearly indicating that the oscillations are no longer present. These results suggest that increasing cell density might accelerate the development of oscillations of the core clock genes via some degree of differentiation, though the kinetics seem to be faster than that of RA induced differentiation. It is interesting to note that in some cases it is impossible to visually discern whether or not an oscillation is present (for example in Figure 5A). These examples serve to highlight the importance of a quantitative and methodical approach to the oscillation detection problem.

## Discussion

Detecting biological oscillations from time-series datasets remains a challenging task even after decades of research. Here we developed an oscillation detection method based on Gaussian Process (GP) regression for learning noisy patterns, combined with Bayesian model selection to distinguish between oscillatory and non-oscillatory datasets. Our method, ODeGP, is designed particularly to model non-stationarity in oscillatory data using the non-stationary kernels introduced recently [36, 37], thereby setting it apart from the few previous GP-based approaches to oscillation detection. Furthermore, the combination of GPs and Bayesian inference has a number of general advantages over existing methods: (1) the non-parametric nature of GPs allows for better learning of noisy, non-stationary and irregularly spaced patterns, (2) technical replicates can easily be incorporated via diagonal terms in the covariance matrix, (3) the learned function and error estimates are naturally generated via analytical expressions of the posterior mean and variance and (4) issues associated with p-values are circumvented by the Bayes factor, which explicitly addresses the likelihood of the observed data under both (oscillatory and non-oscillatory) hypotheses, instead of just the null [32]. Additionally, we also provide a Bonferroni-like multiple hypothesis correction within the Bayesian setting that ODeGP uses [38].

We demonstrated the improved performance of ODeGP compared to eight existing methods using both artificial (simulated) as well as experimental datasets. On the 44 simulated datasets, we used ROC curves and the AUC metric to make the comparisons. Overall, ODeGP worked significantly better (most number of times amongst the top 3 performing methods) than all other methods tested, both for non-stationary as well as stationary symmetric datasets (Tables 2 and 3). As expected however, RAIN consistently outperformed all other methods when the data was asymmetric. Importantly, even when ODeGP did not come out on top for particular datasets, it was still consistently good, as it almost never ranked amongst the 3 worst performing methods. This was in stark comparison to eJTK and Cosinor for example, which often performed very well, but also frequently performed very poorly (Tables 2 and 3). In the case of experimental data where the ground truth is known, we compared the performance of all the methods on low-amplitude oscillations of the *Per1* gene. While most methods incorrectly classified the data as arrhythmic, only ODeGP, eJTK and MetaCycle managed to detect the oscillations (Table 4). However, on calculating the upper bounds of the Bayes factors corresponding to eJTK and MetaCycle’s p-values, we found that these bounds were about two orders of magnitude less than the Bayes Factor generated by ODeGP, thus providing significantly less confidence in the rhythmicity classification. Furthermore, starting from a strongly oscillatory dataset and subsampling the data points, we demonstrated that ODeGP consistently produced higher Bayes factors than other methods. In summary, analysis of both simulated as well as experimental datasets suggests that ODeGP is a more sensitive and reliable oscillation detector in comparison to the existing methods tested here.

Finally, we tested ODeGP’s ability to provide new biological insights in qPCR data on circadian clock genes from pluripotent mESCs. Embryonic stem cells from both mouse [39] and human [43] are known to be deficient in the circadian clock oscillations, even though the genes are expressed in these cells. On directed differentiation using Retinoic Acid, oscillations have been shown to develop on a time scale of about two weeks in these cell types, raising intriguing questions on the role of gradual development of the oscillations [40]. Here we demonstrated that clear oscillations in one core circadian clock gene *Bmal1*, and to a lesser extent in *Per2*, can be induced within 3-4 days by increasing cell density (Figure 5B). Interestingly, we found that these oscillations were prevented from developing in the presence of the inhibitors commonly known as 2i (Figure 5C), thereby suggesting a potential role of density-dependent mESC differentiation in the establishment of the oscillations. Recent experiments have uncovered cell-density effects on strengthening of the clock oscillations [44, 45], potentially via inter-cellular TGF*β* signalling [45]. Our results suggest the intriguing possibility that the kinetics of oscillation development in stem cells could depend on cell-cell signalling during differentiation, and remains an exciting avenue to be further studied in the future.

The oscillatory (alternate hypothesis) versus non-oscillatory (null) classification problem will necessarily involve defining somewhat arbitrary cutoffs. However, based on recent discussions on p-values and interpretation of Bayes Factors as odds ratios, it has been proposed that a p-value of 0.005 is a more sensible cutoff compared to the widely used 0.05 value, corresponding to a Bayes Factor Bound of *∼* 14 [33]. We empirically notice that this guideline seems to be approximately consistent with our own experimental observations. In our case, the real “gold standard” non-oscillatory datasets are the pluripotent mESC datasets (Fig. 5A), where we expect no oscillations to be present even at the single cell level. In these datasets, we consistently find that the Bayes Factors produced by ODeGP are below 14. The interpretation of the unsynchronized (vehicle treated), non-mESC datasets such as in Figure 4A is more challenging. While these datasets are supposed to be non-oscillatory, we observe Bayes Factors that are somewhat higher (16.5 in Fig. 4A and as high as 274 in Fig. S2B). This suggests the presence of weak oscillations, which might be arising from plating of cells and/or addition of fresh media, which are both known to induce a small degree of synchronization between the single cell oscillators. After Dex synchronization however, the oscillations are expected to be stronger, which is correctly being reflected in each case by the much higher Bayes Factors (Fig. 4B,C). Overall, consistent with the recent recommendations in the statistics community, our results suggest that a Bayes Factor cutoff of 14 might be a good choice for classifying oscillatory versus non-oscillatory datasets. In addition, Bayes factors close to 14 could be classified as weak oscillations.

Though not explored extensively, there are a few prior examples of the use of GPs in modeling biological data, for example in identifying differentially expressed genes [36, 46], detecting oscillations [47, 48] and the discovery of spatial patterns in gene expression [49]. The method in [48] uses a principle similar to ours with the comparison of a non-oscillatory GP model to an oscillatory GP model. However, this method does not explicitly incorporate a non-stationary functional form to encode correlations between the data at different time points in its oscillatory model, and also compares its performance only with that of the LS Periodogram (which never falls among the best 3 performing methods in our analysis). GPrank [46] on the other hand uses GPs to model genome-wide time series data. Though it was not created for the specific purpose of detecting oscillations, the workflow followed by this method is similar to ours in terms of the combination of two GP models with Bayesian model selection. The significant difference between GPrank and ODeGP is in the choice of kernel used to define the alternate hypothesis (the RBF kernel is used in GPrank). Our results show that ODeGP almost always outperforms GPrank (Tables 2 and 3), demonstrating the importance of careful choice of the kernel for the specific problem at hand. This point also highlights the flexibility of GPs in modeling various kinds of datasets simply by appropriate choice of kernel functions. Similar to GPrank, the output metric used by ODeGP to make a decision on the presence of oscillations is the Bayes factor, which is a ratio of the marginal likelihoods of the competing models. The marginal likelihood 17 computed during GP regression automatically incorporates a measure of the complexity of the model being considered through the log|*K*| term. A larger log|*K*| for a given model generally results from higher covariance values between the data’s values at different time points, which produce wider confidence bounds in the GP posterior. Wider bounds allow a greater flexibility of functions in the posterior distribution, indicating that the model is more complex. For this reason, we did not additionally penalise the number of hyperparameters of both models when comparing the two (something that is generally done in regression methods which use the BIC/AIC for model selection). Importantly, we found that the Bayes factor was distinctly different between datasets that were known to be oscillatory versus non-oscillatory, thus highlighting the usefulness of this metric in the classification problem.

While our oscillation detection method seems to significantly improve upon currently used methods, there are a number of limitations and potential areas of improvement. As can be seen from Tables 2 and 3, ODeGP is clearly inferior to RAIN in detecting oscillations in asymmetric waveforms. Providing ODeGP an enhanced ability to model asymmetric oscillations represents a clear avenue for improvement, which may be done by identifying new suitable functional forms for the hyperparameters of the non-stationary kernel. The runtime of our method is also significantly higher than that of most of the existing methods we tested (though it is comparable to that of RAIN), due to the hyperparameter optimization routine we used. Furthermore, the multiple hypothesis correction provided with ODeGP is similar to the Bonferroni correction in the frequentist setting [38]. This correction results in many false negatives when there are many hypotheses to be tested, and therefore ODeGP can currently be used only with smaller datasets such as those generated in qPCR, eclosion, egg-laying or feeding experiments. Detecting rhythmic or differentially rhythmic genes from genome-wide datasets is not possible with the current implementation of our method, and remain avenues for future improvement.

## Conclusions

While much work has been done in developing algorithms to detect oscillations in time series datasets, there is clearly room for significant improvement. Our method combining Gaussian processes and Bayesian model selection demonstrates the flexibility of GPs, providing a highly sensitive approach for classifying oscillatory versus non-oscillatory datasets without using p-values. We hope that our results along with the user-friendly ODeGP R package, will spur more careful exploration of these approaches in the future. Applied to new experimental data, ODeGP provides initial evidence for rapid development of circadian clock oscillations upon increasing density of mESCs, and it would be exciting to see in future if these results have broader implications in the context of development and inter-cellular signalling.

## Software Availability

ODeGP is provided as an easy to use R package with this manuscript, and can be downloaded from either https://github.com/shabnamsahay/ODeGP or https://github.com/Shaonlab/ODeGP

## Acknowledgements

S.C. acknowledges funding from SERB (Government of India) under project number SPR/2021/000486 as well as intramural funds from National Center for Biological Sciences–Tata Institute of Fundamental Research (NCBS-TIFR). S.H. acknowledges funding from the National Institute of Health (NIH) National Heart, Lung, and Blood Institute grant no. R01HL158269.

## Author contributions

S.C. conceptualized and designed the study, S.S. developed and implemented the ODeGP algorithm, S.C. and S.A. performed the experiments, S.A. developed the error analysis method for qPCR datasets, S.S. and S.C. wrote the paper with inputs from S.A. and S.H., S.C. and S.H. supervised the work.

