## Supplementary Information for "An improved rhythmicity analysis method using Gaussian Processes detects cell-density dependent circadian oscillations in stem cells"

### Contents

1 qPCR data collection protocol 1

2 Fold-change and error analysis for qPCR datasets 2

3 An overview of Gaussian Process Regression 4

4 Implementation details of ODeGP 6

5 Evaluation against existing methods 12

6 Analysis of model complexity penalisation by MLL 16

7 Application of ODeGP to qPCR datasets 18

### 1 qPCR data collection protocol

For the qPCR experiments to investigate the presence/absence of oscillations, we used the following protocol for Samples 1-4. A frozen vial of mESCs was first thawed and grown on a 100 mm plastic dish coated with gelatin, under pluripotent conditions, till about 60-70% confluency. The cells were then split and grown on a six well plate for about a week (with passaging) under the conditions mentioned in Table 1. Finally, for Dex synchronization and Trizol collection, the cells were transferred to 24 well-plates under the same growth conditions as mentioned in Table 1. To ensure equal cell density at time of Trizol collection, during transfer to the 24-well plates two groups were created - “Day 1” cells were plated into 5 wells of a 24-well plate corresponding to eventual collection time-points 12, 15, 18, 21 and 24 hours, while “Day 2” cells were plated on 4 wells of a separate 24-well plate representing eventual collection time-points 0, 3, 6 and 9 hours. Thus one full 24-hour cycle with 3-hour sampling was represented. For the Day 1 cells, 25000 cells were seeded per well while for the Day 2 cells, 12500 cells were seeded per well. After allowing a few hours for attachment in the 24 well plates, the Day 1 cells were Dex synchronized beginning from 9 am in the following manner: time-point 24 synchronized at 9am, time-point 21 at 12pm, all the way till time-point 12 which was synchronized at 9 pm. Finally, these Day 1 cells were Trizol collected next day at 10 am. Two hours later, beginning from 12 pm, the

Day 2 cells were synchronized every three hours, and finally collected using Trizol at 10 pm. This somewhat elaborate strategy was followed to ensure that at time of collection, all time points had similar cell densities. Assuming an average division time of 12 hours for mESCs, this protocol allowed for 2 doubling times for the Day 1 cells and 3 doubling times for the Day 2 cells. The final expected cell numbers at time of Trizol collection are shown in Table 1. For Samples 5-6, an identical protocol was followed, but with an extra day and an additional collection time, to allow sample RNA collection over 36 hours. Samples 1-4 are considered "low density" since at collection they were about 40-60% confluent while Samples 5-6 are considered "high density" since they were almost completely confluent at time of collection. Note that this high density was achieved only within the 2-3 days that the cells were plated in the 24-well plates, since prior to that the cells were always maintained at less than 60-70% confluency in the 100 mm dish or 6-well plate.

| **Sample number** | **Growth conditions** | **Seeding density** | **Expected cell number at time of Trizol collection** |
|---|---|---|---|
| 1; 24 hrs | Gelatin on plastic, media with LIF | Day 1 cells - 25000<br>Day 2 cells - 12500 | 100000 ("low density") |
| 2; 24 hrs | Fibronectin on glass, media with LIF | Day 1 cells - 25000<br>Day 2 cells - 12500 | 100000 ("low density") |
| 3; 24 hrs | Fibronectin on glass, media with LIF | Day 1 cells - 40000<br>Day 2 cells - 20000 | 160000 ("low density") |
| 4; 24 hrs | Laminin on glass, media with LIF | Day 1 cells - 40000<br>Day 2 cells - 20000 | 160000 ("low density") |
| 5; 36 hrs | Laminin on glass, media with LIF | Day 0 cells - 20000<br>Day 1 cells - 10000<br>Day 2 cells - 5000 | 320000 ("high density") |
| 6; 36 hrs | Laminin on glass, media with LIF + 2i | Day 0 cells - 20000<br>Day 1 cells - 10000<br>Day 2 cells - 5000 | $\sim$ 320000 ("high density") |

Table 1: Description of various samples, seeding densities and substrates used for the qPCR experiments

### 2 Fold-change and error analysis for qPCR datasets

#### Fold change computation

We collected Ct values for a gene of interest from three technical replicates of each sample. For every sample, we also collected three Ct values for the reference gene (i.e. GAPDH). The calculation for fold change is as follows:

- We take the average for three samples ($\langle C_s \rangle$) and three reference samples($\langle C_r \rangle$):

$$\langle C_s \rangle = \frac{1}{3}\sum_{i=1}^{3} C_s^{(i)} \tag{1}$$

$$\langle C_r \rangle = \frac{1}{3}\sum_{i=1}^{3} C_r^{(i)} \tag{2}$$

- We calculate the difference of these two: $\Delta\text{ct} = \langle C_s \rangle - \langle C_r \rangle$.
- $\Delta$ct at time point zero is used as reference $\Delta$ct. We named it $\Delta\text{ct}_{\text{Ref}}$.
- We calculate $\Delta\Delta$ct as the difference between $\Delta$ct and $\Delta\text{ct}_{\text{Ref}}$.
- The fold change is defined as $2^{-\Delta\Delta\text{ct}}$

#### Error propagation

To calculate the error in fold change, we did basic error propagation computation as follows:

- Uncertainty in fold change depends on the error in $\Delta\Delta$ct as follows:

$$\delta_{fc} = \delta_{fc/\Delta\Delta\text{ct}} = \left|\frac{\partial 2^{-\Delta\Delta\text{ct}}}{\partial \Delta\Delta\text{ct}}\right| \delta_{\Delta\Delta\text{ct}} \tag{3}$$

  where, $\delta_{fc/\Delta\Delta\text{ct}}$ is uncertainty in fold change due to uncertainty in $\Delta\Delta$ct and $\delta_{\Delta\Delta\text{ct}}$ is uncertainty in $\Delta\Delta$ct

- $\delta_{\Delta\Delta\text{ct}}$ depends on uncertainty in $\Delta\Delta$ct due to uncertainty in $\Delta$ct ($\delta_{\Delta\Delta\text{ct}/\Delta\text{ct}}$) and uncertainty in $\Delta\Delta$ct due to uncertainty in $\Delta\text{ct}_{\text{Ref}}$ ($\delta_{\Delta\Delta\text{ct}/\Delta\text{ct}_{\text{Ref}}}$) as follows:

$$\delta_{\Delta\Delta\text{ct}} = \sqrt{(\delta_{\Delta\Delta\text{ct}/\Delta\text{ct}})^2 + (\delta_{\Delta\Delta\text{ct}/\Delta\text{ct}_{\text{Ref}}})^2} \tag{4}$$

- $\delta_{\Delta\Delta\text{ct}/\Delta\text{ct}}$ can be calculated as follows:

$$\delta_{\Delta\Delta\text{ct}/\Delta\text{ct}} = \left|\frac{\partial(\Delta\Delta\text{ct})}{\partial \Delta\text{ct}}\right| \delta_{\Delta\text{ct}} = -\Delta\text{ct}_{\text{Ref}} \times \delta_{\Delta\text{ct}} \tag{5}$$

where $\Delta\Delta\text{ct} = \Delta\text{ct} - \Delta\text{ct}_{\text{Ref}}$ and $\delta_{\Delta\text{ct}}$ is the uncertainty in $\Delta\text{ct}$

- Similarly, $\delta_{\Delta\Delta\text{ct}/\Delta\text{ct}_{\text{Ref}}}$ can be calculated as follows:

$$\delta_{\Delta\Delta\text{ct}/\Delta\text{ct}_{\text{Ref}}} = \left|\frac{\partial(\Delta\Delta\text{ct})}{\partial\Delta\text{ct}_{\text{Ref}}}\right|\delta_{\Delta\text{ct}_{\text{Ref}}} = \Delta\text{ct} \times \delta_{\Delta\text{ct}_{\text{Ref}}} \tag{6}$$

where $\delta_{\Delta\text{ct}_{\text{Ref}}}$ is the uncertainty in $\Delta\text{ct}_{\text{Ref}}$

- $\delta_{\Delta\text{ct}}$ can be calculated as follows:

$$\delta_{\Delta\text{ct}} = \sqrt{(\delta_{\Delta\text{ct}/\langle C_s\rangle})^2 + (\delta_{\Delta\text{ct}/\langle C_r\rangle})^2} \tag{7}$$

- $\delta_{\Delta\text{ct}_{\text{Ref}}}$ is just the special case of equation 7, calculated at time point zero.

- $\delta_{\Delta\text{ct}/\langle C_s\rangle}$ can be calculated as follows:

$$\delta_{\Delta\text{ct}/\langle C_s\rangle} = \left|\frac{\partial\Delta\text{ct}}{\partial\langle C_s\rangle}\right|\delta_{\langle C_s\rangle} = -\langle C_r\rangle \times \delta_{\langle C_s\rangle} \tag{8}$$

where $\delta_{\langle C_s\rangle}$ standard error of mean for $C_s(i)$

- $\delta_{\Delta\text{ct}/\langle C_r\rangle}$ can be calculated as follows:

$$\delta_{\Delta\text{ct}/\langle C_r\rangle} = \left|\frac{\partial\Delta\text{ct}}{\partial\langle C_r\rangle}\right|\delta_{\langle C_r\rangle} = -\langle C_s\rangle \times \delta_{\langle C_r\rangle} \tag{9}$$

where $\delta_{\langle C_r\rangle}$ standard error of mean for $C_r(i)$

### 3 An overview of Gaussian Process Regression

Gaussian processes provide a framework for performing non-parametric regression to estimate a target function via Bayesian inference. [1]

A Gaussian process is a collection of random variables, any finite combination of which has a multivariate normal distribution. If each random variable represents the value assumed by the target function $f$ at a particular domain point $x$, then the joint distribution of these random variables corresponds to a distribution over possible target functions.

Let $X^*$ be the vector of domain points $x \in \mathbb{R}$ at which the value of $f$ is to be estimated. Then the prior distribution of $f(X^*)$ (before any data is observed) is given by

$$f(X^*) \sim \mathcal{N}(\mu(X^*), \Sigma_{X^*X^*}) \tag{10}$$

Here, $\mu : \mathbb{R} \to \mathbb{R}$ is a 'mean' function such that $\mu(X^*) = \mathbb{E}[f(X^*)]$. The covariance matrix $\Sigma_{X^*X^*}$ is determined by a positive semi-definite kernel function $K : \mathbb{R}^2 \to \mathbb{R}$ as follows:

$$\Sigma_{ij} = Cov(f(x_i), f(x_j)) = K(x_i, x_j) \text{ for } x_i, x_j \in X^* \tag{11}$$

$K$ thus describes the similarity between values of $f$ at different domain points. The parameters of $K$ are considered to be hyperparameters in the context of the regression. With $\mu$ and $K$ being well-defined on $\mathbb{R}$ and $\mathbb{R}^2$ respectively, $X^*$ can contain any number of $x \in \mathbb{R}$, allowing us to model $f$ at a theoretically infinite number of domain points.

Given a vector of $N$ observed data values $Y$ at corresponding domain points $X$, Bayesian inference can be carried out to obtain a posterior distribution for $f$:

$$f(X^*|Y) \sim \mathcal{N}(\mu^*(X^*), \Sigma^*_{X^*X^*}) \tag{12}$$

$$\text{where } \mu^*(X^*) = \mu(X^*) + \Sigma_{X^*X}\Sigma_{XX}^{-1}(Y - \mu(X))$$

$$\text{and } \Sigma^*_{X^*X^*} = \Sigma_{X^*X^*} - \Sigma_{X^*X}\Sigma_{XX}^{-1}\Sigma_{XX^*}$$

An optimal posterior that best models the given data while avoiding overfitting can be found by maximization of the data likelihood $P(Y|f) = \mathcal{N}(Y|\mu, K)$. The log marginalized form of this likelihood can be expressed as

$$\log P(Y|\mu, K) = -\frac{1}{2}Y^T K_{XX}^{-1} Y - \frac{1}{2}\log |K_{XX}| - \frac{N}{2}\log (2\pi) \tag{13}$$

With the hyperparameters of $K$ being the only variables here, this marginal log likelihood can be maximized by finding optimal values for the hyperparameters using standard techniques.

### 4 Implementation details of ODeGP

#### Pre-processing the time-series data

Let $X$ be a list of $N$ points in the domain at which the data has been observed. In our case, $X$ is a list of times. If $d$ distinct replicates of the data have been obtained for the $N$ time points in $X$, let these be $Y_1, Y_2, \ldots, Y_d$. The average and standard deviation of these $d$ lists at each of the $N$ domain points is calculated to obtain $Y$ and $S$ respectively. This step is not carried out if $Y$ and $S$ are directly supplied as input to the method.

Next, $Y$ is detrended. This is done by performing simple linear regression on the points in $Y$, and setting the new $Y$ as the difference between the initial $Y$ and the values of the resulting best-fit line at the corresponding time points in $X$.

Finally, $Y$ is 0-centered by setting $Y = Y - mean(Y)$. This allows us to set the GP prior distribution's mean as zero, and simplifies further computations required for GP regression.

#### Kernel definitions

The diagonal kernel $K_D$ and the non-stationary kernel $K_{NS}$ are positive semi-definite functions $(R \times R) \to R$ defined as follows:

$$K_D(x, x') = \epsilon^2 \cdot \delta_{xx'} \tag{14}$$

$$\text{where } \delta_{xx'} = \begin{matrix} 0, & x \neq x' \\ 1, & x = x' \end{matrix}$$

$$K_{NS}(x, x') = \sum_{i=1}^{Q} w_i(x) w_i(x') k_{gibbs,i}(x, x') \cos\left(2\pi \left(\frac{x}{\mu_i(x)} - \frac{x'}{\mu_i(x')}\right)\right) + \epsilon_i^2 \cdot \delta_{xx'} \tag{15}$$

We use $K_{NS}$ with $Q = 1$, resulting in the following updated expression:

$$K_{NS}(x, x') = w(x) w(x') k_{gibbs}(x, x') \cos\left(2\pi \left(\frac{x}{\mu(x)} - \frac{x'}{\mu(x')}\right)\right) + \epsilon^2 \cdot \delta_{xx'} \tag{16}$$

where $k_{gibbs}(x, x') = \sqrt{\frac{2l(x)l(x')}{l(x)^2 + l(x')^2}} \exp\left(-\frac{(x - x')^2}{l(x)^2 + l(x')^2}\right)$ , and

$$w(x) = w_{max} - (w_{max} - w_{min})e^{-c_w t}$$

$$l(x) = l_{max} - (l_{max} - l_{min})e^{-c_l t}$$

$$\mu(x) = \mu_{max} - (\mu_{max} - \mu_{min})e^{-c_\mu t}$$

The diagonal kernel has a single hyperparameter $\epsilon$, whereas the non-stationary kernel has ten hyperparameters ($\epsilon$, along with three each for the given functional forms of $w(x), l(x)$ and $\mu(x)$). $\epsilon$ acts as a global noise term in these kernels. The local noise encoded by $S$ is incorporated as follows: for $s_i \in S$, $s_i^2$ is added to the $i$th diagonal entry of the covariance matrix (irrespective of which kernel it is constructed with).

#### The optimization routine

Given the data, optimal hyperparameters for each of $K_D$ and $K_{NS}$ are found through maximization of the marginal log likelihood (MLL). This is the log likelihood of observing the data, given the kernel and the domain points at which the observations are made, i.e. $\log P(Y|K, X)$.

$$\begin{aligned} \text{MLL} &= \log P(Y|K, X) = \log \mathcal{N}(Y|0, K_{XX}) \\ &= -\frac{1}{2}Y^T K_{XX}^{-1} Y - \frac{1}{2}\log |K_{XX}| - \frac{N}{2}\log (2\pi) \end{aligned} \tag{17}$$

where $K_{XX}$ is $K(X, X)$, i.e. the $N \times N$ covariance matrix generated by applying the kernel function $K$ on all pairs of points $(x_i, x_j)$ such that $x_i, x_j \in X$, and then adding $s_i^2$ to $K_{ii}$ for $s_i \in S$.

The DEoptim [2] package, which uses a differential evolution algorithm, is used to carry out this optimization in ODeGP. The DEoptim method is designed to exclusively carry out minimizations, so the negative MLL is passed to this method as the function to be optimized.

The (negative) MLL, as seen from 17, is a function of $X, Y$ and $K$. Since both $X$ and $Y$ are fixed by a given dataset, the optimization is done by searching over different possible $K$, or more specifically different sets of hyperparameters for $K$.

To carry out its optimization, DEoptim requires specification of the upper and lower bounds of the search space for each hyperparameter. For the purpose of defining these bounds, some measures are derived from $X$ and $Y$:

- `X_range`: $max(X) - min(X)$
- `timestep`: $\frac{max(X)-min(X)}{length(X)+1}$
- `Y_range`: $max(Y) - min(Y)$
- `Y_step`: the maximum difference between two consecutive values in the list obtained by arranging the values in $Y$ in ascending order

$K_D$ has a single hyperparameter, $\epsilon$. Since this encodes global noise in ODeGP, we pass the lower and upper bounds for this hyperparameter to DEoptim as 0 and `Y_range` respectively. The same bounds are used for the $\epsilon$ hyperparameter in $K_{NS}$.

The bounds for the remaining nine hyperparameters in $K_{NS}$ are defined in Table 2:

| Hyperparameter | Lower bound | Upper bound |
|---|---|---|
| $w_{max}$ | `Y_step` | `Y_range` |
| $w_{min}$ | 0 | `Y_step` |
| $c_w$ | 0 | `X_range` |
| $l_{max}$ | `timestep` | `X_range` |
| $l_{min}$ | 0 | `timestep` |
| $c_l$ | 0 | `X_range` |
| $\mu_{max}$ | `X_range/2` | `X_range` |
| $\mu_{min}$ | `timestep` | `X_range/2` |
| $c_\mu$ | 0 | `X_range` |

Table 2: List of the hyperparameters and their lower and upper bounds used in optimization of the marginal log likelihood.

These hyperparameters control the variation of $w(x)$, $l(x)$ and $\mu(x)$ through the functional form described in Equation 16, which is introduced in [3]. Their bounds are set as above with the following reasoning:

- $w_{min}$ and $w_{max}$ are respectively the minimum and maximum possible value taken by $w(x)$, which controls the amplitude of the correlation between the GP posterior's values at different time points. Since Y_range is the amplitude of the data as a whole, we take $w_{max} \leq$ Y_range. $w_{min}$ cannot exceed Y_step, else $w(x)$ will not be able to generate correlations with amplitudes comparable to the difference between closely spaced $Y$ values. Y_step thus also acts as a lower bound for $w_{max}$.

- $l_{min}$ and $l_{max}$ are respectively the minimum and maximum possible value taken by $l(x)$, which encodes the lengthscale of correlation between the GP posterior values at different time points. It is clear that $0 \leq l(x) \leq$ X_range. $l_{max}$ cannot be smaller than timestep, else $l(x)$ will not be able to generate correlations between consecutive time points. timestep thus also acts as an upper bound for $l_{min}$.

- $\mu_{min}$ and $\mu_{max}$ are respectively the minimum and maximum possible value taken by $\mu(x)$, which controls the GP posterior's oscillation period at different time points. $\mu_{min} \geq$ the sampling interval of the data (i.e. timestep). $\mu_{max} \leq$ the total time range of the data (X_range), since for time periods > X_range a complete oscillation will not be present in the data. $\mu_{max}$ cannot be smaller than X_range/2, else larger time periods possibly present in the data will not be captured. X_range/2 thus also acts as an upper bound for $\mu_{min}$.

- $c_w$, $c_l$ and $c_\mu$ control the 'curvature' of the functions $w(x)$, $l(x)$ and $\mu(x)$; i.e. the speed with which they respectively approach their respective maximum values [3]. All of these are bounded between 0 and X_range, with the assumption that this curvature should not exceed the total domain range of the data.

The minimization of negative MLL is done with a maximum of 50 iterations for $K_D$, and 200 iterations for $K_{NS}$. Further parameter values used that are specific to DEoptim's implementation can be found in the code for ODeGP.

When complete, the optimization returns the set of hyperparameter values $h$ for the kernel in consideration that results in the minimum attained negative MLL value. Let this be $h_D$ and $h_{NS}$ for $K_D$ and $K_{NS}$ respectively.

#### Model selection with the Bayes factor

Given $X$ and $Y$, $h_D$ and $h_{NS}$, let the corresponding optimal MLL values for $K_D$ and $K_{NS}$ be $\text{MLL}_D$ and $\text{MLL}_{NS}$ respectively.

The Bayes factor is calculated as a ratio of the marginal likelihoods of two competing hypotheses. In our case, it compares the suitability of the two kernels (models) for the given data. Since the MLL values previously calculated are log marginal likelihoods, the Bayes factor is evaluated by applying the exponential function to the difference of $\text{MLL}_{NS}$ and $\text{MLL}_D$.

$$\text{BayesF} = \exp\ (\text{MLL}_{NS} - \text{MLL}_D) \tag{18}$$

The calculated BayesF is returned by the method as output. The larger the resulting value of BayesF, the stronger the support for $K_{NS}$ being a better model for the data. Hence, if this metric exceeds a defined threshold then the data is judged to be oscillatory, else it is not.

#### The posterior distribution and further processing

The GP posterior distribution for kernel $K$ at any given list of $M$ time points $W$ has a mean $\mu$ and covariance matrix $\Sigma$ which can now be calculated as follows:

$$\mu = K_{XW}^T K_{XX}^{-1} Y \tag{19}$$

$$\Sigma = K_{WW} - K_{XW}^T K_{XX}^{-1} K_{XW} \tag{20}$$

where $K_{XW}$ is the $N \times M$ covariance matrix $K(X, W)$, $K_{WW}$ is the $M \times M$ covariance matrix $K(W, W)$ and $K_{XX}$ is the same as defined in 17. Here, while evaluating the kernel function, the hyperparameters of $K$ are fixed as $h$, obtained through optimization of the MLL.

The confidence interval of $\mu$ at time point $w_i \in W$ is evaluated as $2\sqrt{\Sigma_{ii}}$. The strongest detected oscillation period in $\mu$ is obtained using R's `spectrum` function from the `stats` library.

#### Correction to the Bayes Factor for multiple-hypothesis testing

While ODeGP is primarily intended for detecting oscillations in single trajectories, a multiple hypothesis testing correction in the Bayesian setting is also provided in case a dataset has multiple trajectories that need to be simultaneously analyzed for the presence of rhythms. Here we provide a brief explanation of the correction factor, and how it compares with the frequentist Bonferroni approach of controlling the Family-Wise Error Rate.

As discussed above, ODeGP computes the Bayes factor as the ratio of the marginal likelihoods of the alternate (oscillatory; $H_A$) versus null (not-oscillatory; $H_0$) hypotheses. Using Bayes theorem, it is easy to show that:

$$\frac{\pi(H_A|Y)}{\pi(H_0|Y)} = \frac{\pi(H_A)}{\pi(H_0)} \frac{P(Y|H_A)}{P(Y|H_0)}, \tag{21}$$

where the left hand side represents the posterior odds, and the right hand side is the product of the prior odds and the Bayes Factor. When using just the Bayes factor to classify a waveform as oscillatory or not, ODeGP implicitly assumes that the prior odds is 1, i.e. $\pi(H_A) = \pi(H_0)$. This results in the posterior odds being identical to the Bayes factor. In this setting, multiple hypothesis corrections can be carried out by suitably modifying the prior odds to generate a posterior odds that is less than the Bayes factor [4], as described below.

Assume there are $k$ trajectories that are to be tested for oscillations, with corresponding null and alternate hypotheses $H_{0i}$, $H_{Ai}$ respectively, where $i = 1, ..., k$. Suppose that initially $\pi(H_{0i})$ is set to 0.5 for all $i$, implying that the assessor is truly objective about the prior probability of the hypotheses being true or false. Given independent null hypotheses, the joint probability of all the null hypotheses being true becomes $0.5^k$, which is possibly much smaller than desired. In this case, one could begin with assigning a value for the joint probability $\Pi_0 = \prod_{i=1}^{k} \pi(H_{0i})$, for instance 0.5, and then modifying the $\pi(H_{0i})$ accordingly. If there is no preference for any particular $H_{0i}$, then the revised $\pi(H_{0i})$ becomes equal to $\Pi_0^{1/k}$. In this case, the prior odds for each trajectory is no longer 1 and the posterior odds in Equation. 21 gets modified to:

$$\frac{\pi(H_{Ai}|Y)}{\pi(H_{0i}|Y)} = \left(\frac{1-\Pi_0^{1/k}}{\Pi_0^{1/k}}\right)\frac{P(Y|H_{Ai})}{P(Y|H_{0i})}. \tag{22}$$

Note that the Bayes factor for each hypothesis, $\frac{P(Y|H_{Ai})}{P(Y|H_{0i})}$, is calculated independently as explained previously. ODeGP reports this prior odds as a correction term to the Bayes factor, with a default value $\Pi_0 = 0.5$. It can be easily demonstrated (see [4] for the derivation) that under certain conditions ($k \to \infty$ and 1/Bayes factor $\to 0$), this method of multiple hypothesis correction is similar to the frequentist Bonferroni correction, since the modified posterior probability $\pi(H_{Ai}|Y)$ is scaled by a factor approximately equal to $k$.

### 5 Evaluation against existing methods

#### Generating simulated datasets

Simulated waves were generated as sets of 3 replicates, with a duration of 48 hours and an interval of 3 hours between consecutive observations (when no data was missing). Each such set of 3 replicates was produced as follows: a base waveform was generated at the designated timepoints - let these times be contained in a vector $T$, the corresponding waveform values be contained in a vector $D$, and the length of $T$ = length of $D = N$. Taking the standard deviation of $D$ to be $\sigma_D$, each replicate was generated by adding to $D$, a vector of $N$ independent samples drawn from $U(-\sigma_D, \sigma_D)$.

The base waveform $D$ is generated by applying a function $f(t)$ to the times in $T$ (see Tables 3, 4, 5 and 6 for details). Following this, a fraction $\gamma$ of the times in $T$ are randomly chosen, and these times along with the corresponding values in $D$ are deleted from $T$ and $D$ respectively.

Construction of an ROC curve requires a dataset consisting of both oscillatory and non-oscillatory waves. Each such dataset was generated as a collection of 500 oscillatory waves and 500 non-oscillatory waves, where each wave is a set of 3 replicates produced from a defined base waveform as described above.

For oscillatory waves, the definitions of $f$ depend on the type of waves being simulated, and are provided in Tables 3, 4, 5 and 6 along with the values of $\gamma$ used in different cases. For non-oscillatory waveforms, $f$ was defined as $f(t) = \mathcal{N}(0, \sigma)$, and $\gamma =$ the $\gamma$ being used to generate the 500 oscillatory waves comprising the other half of that particular dataset of 1000.

| **Stationary symmetric waves** | |
|---|---|
| $\mathbf{f(t)}$ | $A_1 \sin(2\pi t/\tau_1) + A_2 \sin(2\pi t/\tau_2) + \mathcal{N}(0, \sigma)$ |
| Dataset no. | Parameter values |
| 1,2,3,4 | $A_1 = 3, \tau_1 = 22, A_2 = 3, \tau_2 = 28$<br>$(\sigma, \gamma) = (0.1, 0.0)$ or $(0.1, 0.5)$ or $(1, 0.0)$ or $(1, 0.5)$ |
| 5,6,7,8 | $A_1 = 3, \tau_1 = 18, A_2 = 3, \tau_2 = 26$<br>$(\sigma, \gamma) = (0.1, 0.0)$ or $(0.1, 0.5)$ or $(1, 0.0)$ or $(1, 0.5)$ |
| 9 | $A_1 = 5, \tau_1 = 24, A_2 = 0, \tau_2 = 24$<br>$(\sigma, \gamma) = (1, 0.5)$ |

Table 3: Functional forms and parameter values used to generate simulated stationary symmetric datasets.

| **Stationary asymmetric waves** | |
|---|---|
| $\mathbf{f(t)}$ | $A \cdot \{t/\tau\} + \mathcal{N}(0, \sigma)$<br>where $\{\}$ represents the fractional part function |
| Dataset no. | Parameter values |
| 10,11,12,13 | $A = 5, \tau = 18$<br>$(\sigma, \gamma) = (0.1, 0.0)$ or $(0.1, 0.5)$ or $(1, 0.0)$ or $(1, 0.5)$ |
| 14,15,16,17 | $A = 5, \tau = 26$<br>$(\sigma, \gamma) = (0.1, 0.0)$ or $(0.1, 0.5)$ or $(1, 0.0)$ or $(1, 0.5)$ |

Table 4: Functional forms and parameter values used to generate simulated stationary asymmetric datasets.

| **Non-stationary symmetric waves** | |
|---|---|
| $\mathbf{f(t)}$ (type 1 - monotonically decreasing time period) | $A \cdot \sin(\frac{2\pi t}{1+\lvert t-\tau \rvert}) + \mathcal{N}(0, \sigma)$<br>where $\tau \geq 48$ |
| Dataset no. | Parameter values |
| 18,19,20,21 | $A = 5, \tau = 72$<br>$(\sigma, \gamma) = (0.1, 0.0)$ or $(0.1, 0.5)$ or $(1, 0.0)$ or $(1, 0.5)$ |
| 22,23,24,25 | $A = 5, \tau = 60$<br>$(\sigma, \gamma) = (0.1, 0.0)$ or $(0.1, 0.5)$ or $(1, 0.0)$ or $(1, 0.5)$ |
| 26,27,28,29 | $A = 5, \tau = 48$<br>$(\sigma, \gamma) = (0.1, 0.0)$ or $(0.1, 0.5)$ or $(1, 0.0)$ or $(1, 0.5)$ |
| $\mathbf{f(t)}$ (type 2 - randomly varying time period) | At $t = 0$ OR at the completion of each time period, a new time period $\tau$ is sampled from $\mathcal{N}(\mu, \sigma)$. If the total time elapsed upto the start of this new oscillation is $\phi$, then $f(t) = A \cdot \sin(\frac{2\pi(t-\phi)}{\tau}) + \mathcal{N}(0, \sigma)$ for $\phi \leq t < \phi + \tau$ |
| Dataset no. | Parameter values |
| 30,31,32 | $\mu = 24, \sigma = 1.33, A = 1.5$<br>$(\sigma, \gamma) = (1, 0.0)$ or $(1, 0.5)$ or $(0.5, 0.5)$ |

Table 5: Functional forms and parameter values used to generate simulated non-stationary symmetric datasets.

| Non-stationary asymmetric waves | |
|---|---|
| $\mathbf{f(t)}$ | At $t = 0$ OR at the completion of each time period, a new time period $\tau$ is sampled from $U(\tau_1, \tau_2)$. If the total time elapsed upto the start of this new oscillation is $\phi$, then $f(t) = A \cdot \{\frac{t-\phi}{\tau}\} + \mathcal{N}(0, \sigma)$ for $\phi \leq t < \phi + \tau$, where $\{\}$ represents the fractional part function |
| Dataset no. | Parameter values |
| 33,34,35,36 | $\tau_1 = 12, \tau_2 = 30, A = 5$<br>$(\sigma, \gamma) = (0.1, 0.0)$ or $(0.1, 0.5)$ or $(1, 0.0)$ or $(1, 0.5)$ |
| 37,38,39,40 | $\tau_1 = 12, \tau_2 = 21, A = 5$<br>$(\sigma, \gamma) = (0.1, 0.0)$ or $(0.1, 0.5)$ or $(1, 0.0)$ or $(1, 0.5)$ |
| 41,42,43,44 | $\tau_1 = 21, \tau_2 = 30, A = 5$<br>$(\sigma, \gamma) = (0.1, 0.0)$ or $(0.1, 0.5)$ or $(1, 0.0)$ or $(1, 0.5)$ |

Table 6: Functional forms and parameter values used to generate simulated non-stationary asymmetric datasets.

### Specification of parameters for running existing methods

Let $X$ be the list of time points at which the data has been observed. We have earlier defined `X_range` $= max(X) - min(X)$. Considering that the individual elements of a list are accessed by 1-indexing in R, we additionally define `delta_t` $= X[2] - X[1]$.

MetaCycle's [5] `meta2d` function requires the minimum and maximum period length of interest to be passed as arguments. When running MetaCycle on our simulated datasets, these were respectively specified as

- `minper` = 2*`delta_t`
- `maxper` = `X_range`

RAIN's [6] `rain` function requires the period to test for and/or the range of periods to test for to be passed as arguments. These were respectively specified during testing as:

- `period` = `X_range`/2
- `period.delta` = ( `X_range`/2)− `delta_t`

For all methods tested that accept the raw data (without merging of replicates) as input, the replicates were passed to the method in an appropriate format.

#### AUC values of tested methods on simulated datasets

The numbers assigned to simulated datasets in the previous section describing their generation are used here. AUC values for the simulated stationary datasets no. 6 and 12 are shown in Figure 2 of the main text. The values for the simulated non-stationary datasets no. 19, 32 and 33 are shown in Figure 3 of the main text.

| Method | AUC value for dataset no. | | | | | | | | | |
|---|---|---|---|---|---|---|---|---|---|---|
| | 1 | 2 | 3 | 4 | 5 | 7 | 8 | 9 | 10 | 11 |
| Cosinor | 1.000 | 0.978 | 0.999 | 0.960 | 0.832 | 0.796 | 0.632 | 0.987 | 0.608 | 0.518 |
| eJTK | 1.000 | 0.954 | 0.999 | 0.930 | 0.766 | 0.729 | 0.586 | 0.968 | 0.324 | 0.444 |
| GPrank | 0.974 | 0.691 | 0.943 | 0.672 | 0.955 | 0.922 | 0.626 | 0.679 | 0.763 | 0.580 |
| JTK_Cycle | 0.999 | 0.874 | 0.994 | 0.836 | 0.955 | 0.931 | 0.666 | 0.897 | 0.959 | 0.720 |
| LSP | 0.998 | 0.811 | 0.988 | 0.754 | 0.942 | 0.898 | 0.545 | 0.867 | 0.969 | 0.680 |
| Metacycle | 0.999 | 0.873 | 0.995 | 0.829 | 0.962 | 0.937 | 0.642 | 0.905 | 0.972 | 0.725 |
| ODeGP | 0.983 | 0.829 | 0.963 | 0.782 | 0.965 | 0.922 | 0.672 | 0.780 | 0.723 | 0.629 |
| SM Kernel | 0.999 | 0.911 | 0.996 | 0.878 | 0.996 | 0.982 | 0.786 | 0.907 | 0.855 | 0.698 |
| RAIN | 0.995 | 0.847 | 0.985 | 0.808 | 0.985 | 0.967 | 0.709 | 0.861 | 0.999 | 0.828 |

| Method | AUC value for dataset no. | | | | | | | | | |
|---|---|---|---|---|---|---|---|---|---|---|
| | 13 | 14 | 15 | 16 | 17 | 18 | 20 | 21 | 22 | 23 |
| Cosinor | 0.516 | 0.968 | 0.820 | 0.884 | 0.712 | 0.884 | 0.890 | 0.705 | 0.927 | 0.742 |
| eJTK | 0.468 | 0.995 | 0.834 | 0.910 | 0.720 | 0.832 | 0.834 | 0.670 | 0.937 | 0.723 |
| GPrank | 0.550 | 0.899 | 0.678 | 0.766 | 0.591 | 0.962 | 0.932 | 0.695 | 0.733 | 0.593 |
| JTK_Cycle | 0.604 | 0.978 | 0.777 | 0.849 | 0.644 | 0.818 | 0.772 | 0.675 | 0.678 | 0.572 |
| LSP | 0.543 | 0.967 | 0.686 | 0.751 | 0.540 | 0.573 | 0.542 | 0.583 | 0.405 | 0.572 |
| Metacycle | 0.592 | 0.982 | 0.769 | 0.829 | 0.621 | 0.760 | 0.710 | 0.660 | 0.584 | 0.580 |
| ODeGP | 0.559 | 0.889 | 0.736 | 0.653 | 0.586 | 0.967 | 0.919 | 0.719 | 0.506 | 0.617 |
| SM Kernel | 0.608 | 0.946 | 0.780 | 0.812 | 0.657 | 0.991 | 0.966 | 0.785 | 0.816 | 0.665 |
| RAIN | 0.615 | 0.994 | 0.785 | 0.866 | 0.628 | 0.756 | 0.746 | 0.626 | 0.650 | 0.532 |

| Method | AUC value for dataset no. | | | | | | | | | |
|---|---|---|---|---|---|---|---|---|---|---|
| | 24 | 25 | 26 | 27 | 28 | 29 | 30 | 31 | 34 | 35 |
| Cosinor | 0.911 | 0.727 | 0.619 | 0.515 | 0.620 | 0.515 | 0.970 | 0.812 | 0.684 | 0.744 |
| eJTK | 0.925 | 0.717 | 0.405 | 0.450 | 0.413 | 0.461 | 0.962 | 0.790 | 0.693 | 0.723 |
| GPrank | 0.721 | 0.587 | 0.661 | 0.562 | 0.632 | 0.574 | 0.783 | 0.842 | 0.660 | 0.738 |
| JTK_Cycle | 0.660 | 0.565 | 0.777 | 0.634 | 0.744 | 0.635 | 0.898 | 0.903 | 0.717 | 0.772 |
| LSP | 0.388 | 0.557 | 0.558 | 0.579 | 0.514 | 0.540 | 0.849 | 0.803 | 0.631 | 0.659 |
| Metacycle | 0.557 | 0.569 | 0.719 | 0.630 | 0.675 | 0.617 | 0.891 | 0.883 | 0.705 | 0.745 |
| ODeGP | 0.493 | 0.599 | 0.574 | 0.583 | 0.566 | 0.593 | 0.689 | 0.763 | 0.679 | 0.614 |
| SM Kernel | 0.790 | 0.660 | 0.787 | 0.652 | 0.752 | 0.634 | 0.909 | 0.887 | 0.733 | 0.766 |
| RAIN | 0.644 | 0.533 | 0.801 | 0.615 | 0.758 | 0.610 | 0.907 | 0.952 | 0.747 | 0.806 |

| Method | AUC value for dataset no. | | | | | | | | |
|---|---|---|---|---|---|---|---|---|---|
| | 36 | 37 | 38 | 39 | 40 | 41 | 42 | 43 | 44 |
| Cosinor | 0.628 | 0.527 | 0.511 | 0.516 | 0.518 | 0.933 | 0.776 | 0.843 | 0.683 |
| eJTK | 0.630 | 0.433 | 0.496 | 0.442 | 0.491 | 0.950 | 0.796 | 0.852 | 0.688 |
| GPrank | 0.594 | 0.788 | 0.626 | 0.700 | 0.597 | 0.901 | 0.686 | 0.768 | 0.613 |
| JTK_Cycle | 0.614 | 0.876 | 0.678 | 0.742 | 0.604 | 0.958 | 0.746 | 0.813 | 0.641 |
| LSP | 0.526 | 0.823 | 0.636 | 0.668 | 0.571 | 0.944 | 0.630 | 0.720 | 0.526 |
| Metacycle | 0.594 | 0.874 | 0.682 | 0.727 | 0.602 | 0.963 | 0.730 | 0.794 | 0.618 |
| ODeGP | 0.586 | 0.646 | 0.639 | 0.566 | 0.575 | 0.850 | 0.702 | 0.627 | 0.586 |
| SM Kernel | 0.621 | 0.787 | 0.647 | 0.673 | 0.586 | 0.934 | 0.751 | 0.808 | 0.644 |
| RAIN | 0.617 | 0.929 | 0.739 | 0.778 | 0.623 | 0.985 | 0.776 | 0.839 | 0.637 |

### 6 Analysis of model complexity penalisation by MLL

The expression for $K_{NS}$ in 15 contains a variable $Q$, which denotes the number of components [7] of $K_{NS}$. Every increase of 1 in $Q$ adds 9 new hyperparameters to the defined $K_{NS}$. We show that increasing $Q$ (and thus the number of hyperparameters) does not cause overfitting of the data in the posterior generated by $K_{NS}$, and that the MLL does penalise the more 'complex' resulting model represented by $K_{NS}$.

Non-oscillatory data was simulated by sampling from $\mathcal{N}(0,1)$ at 3 hour intervals for a duration of 48 hours, and generating three replicates from this base waveform as described above. ODeGP was run on this data with multiple values of $Q$. The datafit and complexity term of the MLL for $K_{NS}$ were respectively calculated as $-0.5Y^T K_{XX}^{-1} Y$ and $-0.5\log K_{XX}$, as in 17.

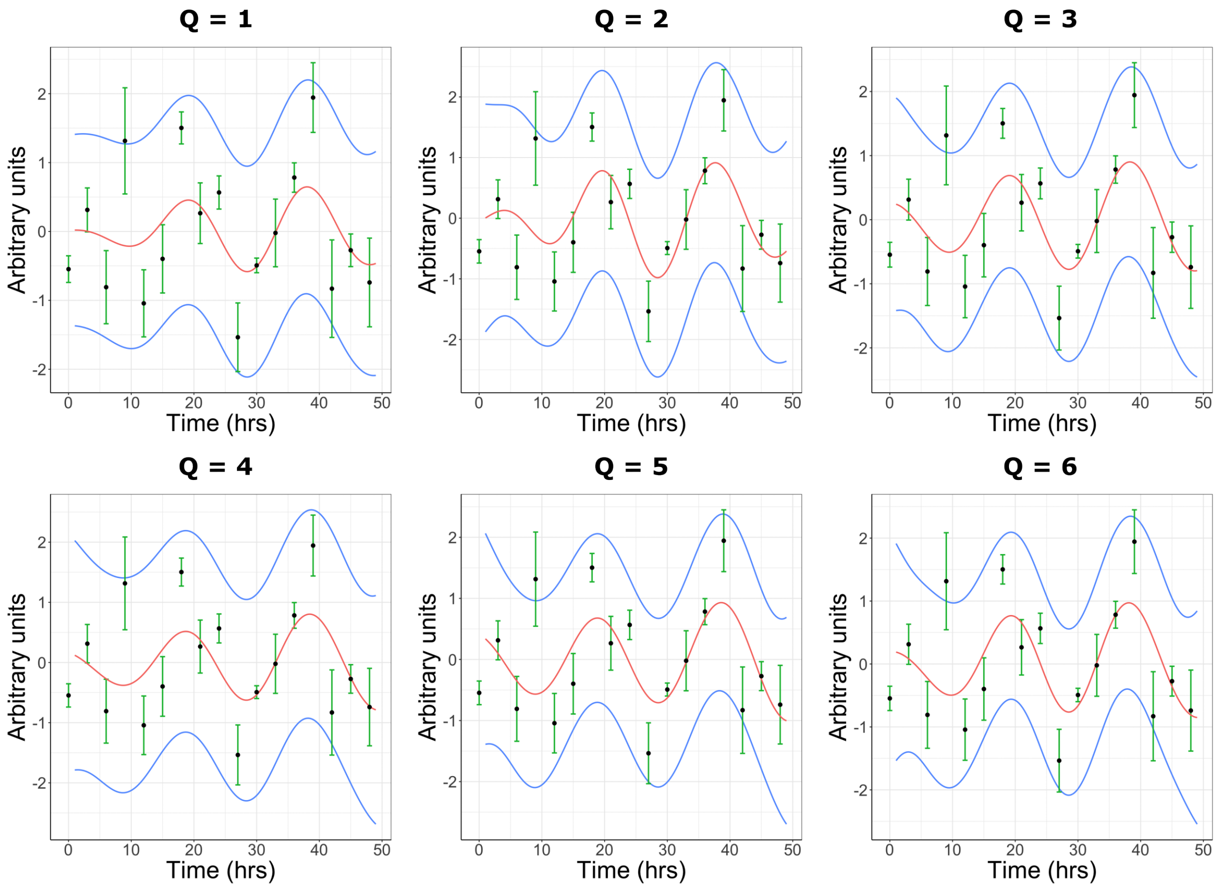


Figure 1: Posterior distributions of the non-stationary kernel with varying number of components $Q$, obtained by application of ODeGP on simulated non-oscillatory data.

| Number of components (Q) used in $K_{NS}$ | Total number of hyperparameters | Bayes factor | Datafit term ($K_{NS}$) | Complexity term ($K_{NS}$) |
|---|---|---|---|---|
| 1 | 10 | 2.2397 | -7.9760 | 1.6739 |
| 2 | 19 | 1.1253 | -5.7758 | -1.2145 |
| 3 | 28 | 0.5657 | -7.5851 | -0.0930 |
| 4 | 37 | 0.1715 | -6.6950 | -2.1765 |
| 5 | 46 | 0.1056 | -8.1752 | -1.1811 |
| 6 | 55 | 0.0438 | -8.3841 | -1.8528 |

Table 7: Comparison of the Bayes factor and individual terms of the MLL for $K_{NS}$, obtained by testing ODeGP on the non-oscillatory data shown in 1 with different values of $Q$ in $K_{NS}$. A higher value of the datafit term indicates a closer fit of the data by the $K_{NS}$ posterior mean, and a higher value of the complexity term indicates a less complex (and more favourable) model generated by $K_{NS}$.

Figure 1 demonstrates that increasing the number of hyperparameters used in the non-stationary kernel does not enforce a closer fitting of the data by the posterior mean. This is also reflected in the datafit term of $K_{NS}$ in Table 7. The complexity term worsens as well (though not monotonically) and there is a clear trend of reduction in the Bayes factor. Overall, a larger $Q$ leads to a stronger indication of $K_D$ representing a better model for the data.

### 7 Application of ODeGP to qPCR datasets

To understand how well ODeGP works on existing qPCR datasets, we analyzed gene expression profiles reported in a prior work [8]. While the expression patterns and analysis of *Rev-ERBβ* and *Per1* were shown in the main text Figure 4, here we present analysis of some of the other reported genes, *Per2* and *Npas2* along with the unsynchronized data for *Per1*. It is clear that between the Dex and vehicle treated datasets, the Bayes factor is always separated by an order of magnitude or more, demonstrating the power of ODeGP in classifying oscillatory datasets.

Similar to the analysis of the synchronized *Per1* data shown in Table 4 of the main text, we also evaluated the Bayes factor bounds produced by each method for the unsynchronized Vehicle-treated *Per1* data in Figure 2C. The results of this are shown in Table 8. Barring MetaCycle and RAIN, all methods correctly classify the data as non-oscillatory with p-values greater than 0.05. The Bayes factor bounds obtained for the remaining methods are all sufficiently close to 1, strongly indicating that the data does not contain oscillations. ODeGP's non-stationary kernel posterior for this data in Figure 2C shows very wide confidence intervals, confirming that this kernel is not a good model for it.

| Method | Metric type | Metric value | Bayes factor (bound) |
|---|---|---|---|
| Cosinor | Q-value | 0.50131 | 1.0000 |
| eJTK | p-value | 0.99289 | 1.0000 |
| GPrank | log of Bayes factor | 0.09877 | 1.1038 |
| JTK_Cycle | p-value | 0.09852 | 1.6112 |
| LSP | p-value | 0.11609 | 1.4716 |
| MetaCycle | p-value | 0.01397 | 6.1659 |
| ODeGP | Bayes factor | 1.1313 | 1.1313 |
| SM Kernel | Bayes factor | 1.3785 | 1.3785 |
| RAIN | p-value | 0.00617 | 11.7184 |

Table 8: Comparison of output metrics of all methods tested on the Vehicle-treated *Per1* data shown in Figure 2C. p-values/Q-values were converted to corresponding Bayes factor bound [9] values where applicable to allow comparison with the Bayes factor returned by ODeGP.

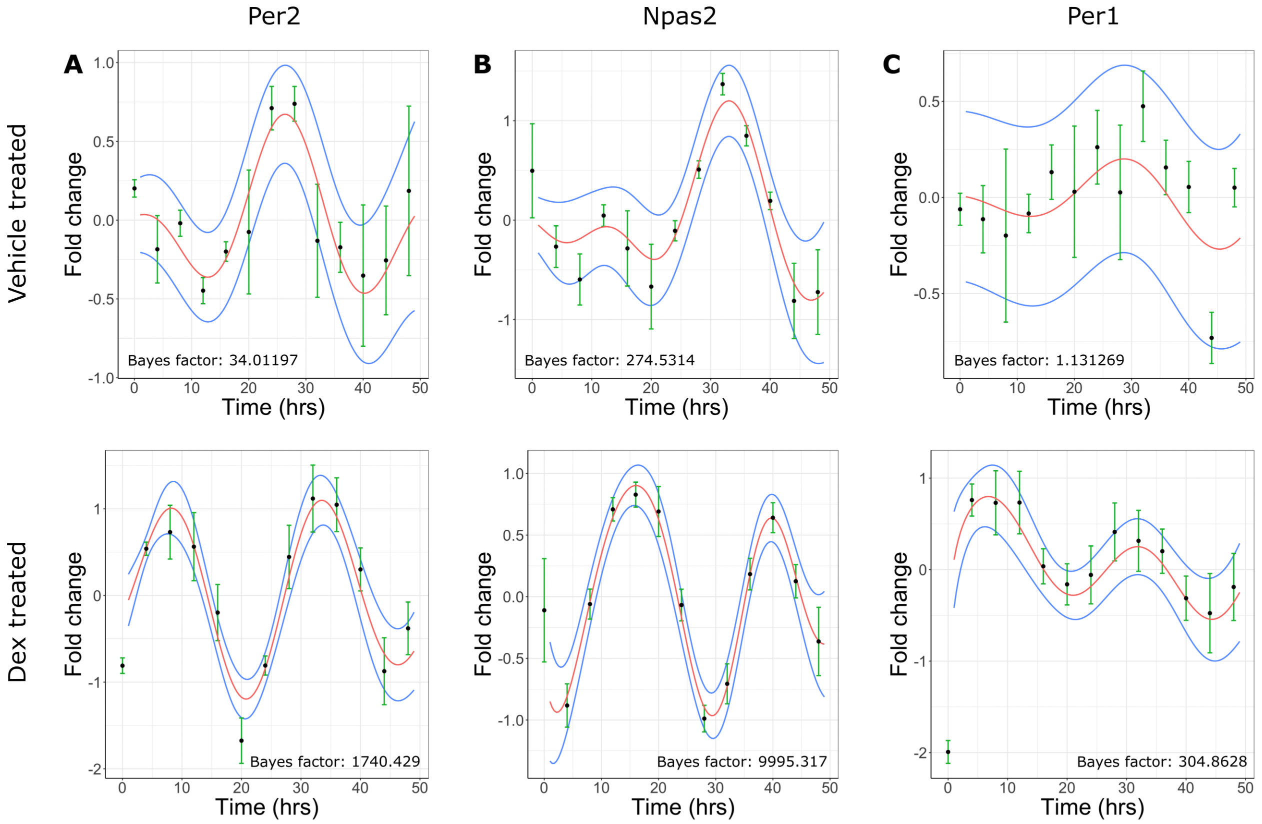


Figure 2: Application of ODeGP on *Per2*, *NPas2* and *Per1* expression data measured using qPCR [8]. (A) The posterior distributions (mean in red, standard deviation in blue) obtained by defining a GP prior using the non-stationary kernel, and performing GP regression on the relative gene expression of *Per2* after treatment with DMSO (above) or Dexamethasone (below), measured at 4 hour intervals over a 48 hour duration and averaged over 3 biological replicates. (B) The GP posteriors of the non-stationary kernel applied to relative gene expression of *Npas2* when treated with DMSO (above) or Dexamethasone (below), where the posterior mean and standard deviation are in red, blue respectively. (C) The GP posteriors of the non-stationary kernel applied to relative gene expression of *Per1* when treated with DMSO (above) or Dexamethasone (below), where the posterior mean and standard deviation are in red, blue respectively.

### References

[1] Carl Edward Rasmussen and Christopher K. I. Williams. *Gaussian Processes for Machine Learning*. MIT Press, November 2005. Google-Books-ID: Tr34DwAAQBAJ.

[2] Katharine Mullen, David Ardia, David Gil, Donald Windover, and James Cline. DEoptim: An R package for global optimization by Differential Evolution. *Journal of Statistical Software*, 40(6):1–26, 2011.

[3] Markus Heinonen, Olivier Guipaud, Fabien Milliat, Valérie Buard, Béatrice Micheau, Georges Tarlet, Marc Benderitter, Farida Zehraoui, and Florence d'Alché Buc. Detecting time periods of differential gene expression using gaussian processes: an application to en-

dothelial cells exposed to radiotherapy dose fraction. *Bioinformatics*, 31(5):728–735, March 2015.

[4] Peter H. Westfall, Wesley O. Johnson, and Jessica M. Utts. A Bayesian Perspective on the Bonferroni Adjustment. *Biometrika*, 84(2):419–427, 1997. Publisher: [Oxford University Press, Biometrika Trust].

[5] Gang Wu, Ron C. Anafi, Michael E. Hughes, Karl Kornacker, and John B. Hogenesch. MetaCycle: an integrated R package to evaluate periodicity in large scale data. *Bioinformatics*, 32(21):3351–3353, 07 2016.

[6] Paul F. Thaben and Pål O. Westermark. Detecting rhythms in time series with RAIN. *Journal of Biological Rhythms*, 29(6):391–400, October 2014.

[7] Sami Remes, Markus Heinonen, and Samuel Kaski. Non-stationary spectral kernels. 2017.

[8] Alex Y.-L. So, Teresita U. Bernal, Marlisa L. Pillsbury, Keith R. Yamamoto, and Brian J. Feldman. Glucocorticoid regulation of the circadian clock modulates glucose homeostasis. *Proceedings of the National Academy of Sciences of the United States of America*, 106(41):17582–17587, October 2009.

[9] Daniel J. Benjamin and James O. Berger. Three recommendations for improving the use of p-values. *The American Statistician*, 73:186–191, 2019.
